# The Notch ligand Jagged1 plays a dual role in cochlear hair cell regeneration

**DOI:** 10.1101/2025.03.02.640998

**Authors:** Xiao-Jun Li, Charles Morgan, Lin Li, Wan-Yu Zhang, Elena Chrysostomou, Angelika Doetzlhofer

**Author notes:** CORRESPONDENCE ADDRESSED TO: Angelika Doetzlhofer, Ph.D., The Solomon H. Snyder Department of Neuroscience, Johns Hopkins School of Medicine, 855 N. Wolfe Street, Rangos Building, Room 433, Baltimore, MD 21205, Xiaojun Li, Ph.D., Frontier Institute of Science and Technology, Xi’an Jiaotong University, No.28 Xianning West Road, Xi’an, Shaanxi 710049.

## Abstract

Hair cells (HCs) within the inner ear cochlea are specialized mechanoreceptors required for hearing. Cochlear HCs are not regenerated in mammals, and their loss is a leading cause of deafness in humans. Cochlear supporting cells (SCs) in newborn mice have the capacity to regenerate HCs, but persistent Notch signaling, presumably activated by SC-specific Notch ligand Jagged1 (JAG1), prevents SCs from converting into HCs. Here, employing an organoid platform, we show that while JAG1 participates in HC-fate repression, JAG1’s primary function is to preserve the “progenitor-like characteristics” of cochlear SCs. Transcriptomic and mechanistic studies reveal that JAG1/Notch signaling maintains the expression of progenitor and metabolic genes in cochlear SCs and sustains pro-growth pathways, including PI3K-Akt-mTOR signaling, a function that is mediated by Notch1 and Notch2. Finally, we show that JAG1/Notch signaling stimulation with JAG1-Fc peptide enhances the HC-forming capacity of cochlear SCs undergoing maturation in cochlear explants and *in vivo*.

## INTRODUCTION

Notch signaling is an evolutionary conserved signaling pathway that, through juxtracrine interactions of its transmembrane receptors and ligands, controls key aspects of vertebrate development (reviewed in^1^). Mammals express four Notch receptors (Notch1-4) and five canonical Notch ligands, Delta-like 1, 3, 4 (DLL1, 3, 4) and Jagged1, 2 (JAG1, 2). Upon ligand binding, the Notch receptor protein undergoes a conformational change, triggering proteolytic cleavages by ADAM and γ-secretase enzymes, which frees the Notch intracellular domain (NICD) to translocate into the nucleus. In the nucleus, NICD binds to the RBPJ-MAML complex to drive the expression of Notch target genes (reviewed in^2^).

Notch signaling has two contrasting functions during the development of the auditory sensory organ. Low Notch receptor signaling strength is sufficient to specify and maintain the pool of progenitors (termed pro-sensory cells) from which two types of mechano-sensory hair cells (HCs), termed inner and outer HCs and several subtypes of surrounding supporting cells (SCs) derive. By contrast, high Notch receptor signaling strength is required to repress HC-fate and limit the number of HCs that are produced during differentiation (reviewed in^3^). These two distinct modes of Notch receptor signaling rely on different sets of Notch ligands. The Notch ligand JAG1, which in the murine cochlea is initially expressed at the medial border of the pro-sensory domain and later throughout the pro-sensory domain, elicits low levels of Notch1 receptor signaling^4,5^, which is critical to maintaining the expression of SOX2^6^, a transcription factor essential for pro-sensory cell fate specification^7^. In addition, JAG1/Notch signaling positively regulates the expression of the transcriptional repressors HEY1^8,9^ and HES1^10^, which help to maintain pro-sensory cells in an undifferentiated state. Highlighting JAG1’s critical role in maintaining a pro-sensory cell fate, early otic deletion of *Jag1* results in a near-complete loss of outer HCs and surrounding SCs ^6,11^. By contrast, the Notch ligands DLL1 and JAG2 are essential for HC-fate repression. DLL1 and JAG2 are expressed by nascent HCs and are capable of activating high levels of Notch1 receptor signaling^4,5^, leading to the induction of HES5^12–14^ in neighboring pro-sensory cells. Together with other members of HES/HEY family of transcriptional repressors HES5^15,16^ antagonizes the function of ATOH1, a transcription factor essential for HC-fate determination^17^, limiting these pro-sensory cells to an SC fate. Consistent with their HC-repressive role, early otic deletion of *Dll1* and or *Jag2* results in supernumerary HCs^11,18,19^. Similarly, early otic deletion of *Notch1* results in massive overproduction of HCs, indicating that the Notch1 receptor is critical for HC-fate repression but dispensable for Notch signaling’s earlier function in pro-sensory cell maintenance^19^.

In mice, cochlear HCs and SCs are formed between embryonic day (E) 14.5 and E18.5, after which they undergo a 2-week long maturation process that ends with the onset of hearing at around postnatal day 13 (P13)^20^. During the early phase of postnatal development (P0-P3), Notch signaling is highly active in cochlear SCs, and inhibition of Notch signaling using a small molecule γ-secretase inhibitor (GSI) is sufficient to convert SCs into HCs^21^. Even in the absence of HCs, the presence of GSI triggers massive conversion of SCs into HCs, indicating that Notch signaling remains active in cochlear SCs following HC loss^22,23^. At later stages (P5 and beyond), the ability of murine cochlear SCs to form HCs in response to GSI treatment sharply declines^24^; nevertheless, GSI drug treatment has been reported to yield new HCs and improve hearing after acoustic trauma in adult mice^23^.

A candidate for preventing SC-to-HC conversion in the absence of HCs is the Notch ligand JAG1, which is expressed in both developing and mature cochlear SCs^25^. Recent studies in mice revealed that conditional deletion of *Jag1* in cochlear SCs at early postnatal stages leads to hearing deficits, which is accompanied by the loss of Hensen cells (SC subtype)^26^ and improper development of inner HC stereocilia^27^, highlighting a critical role for JAG1 in the maturation and maintenance of cochlear SCs and HCs. However, JAG1’s role in SC-mediated HC formation/regeneration is poorly understood.

Using cochlear epithelial-derived organoid cultures, we show that Notch ligand JAG1 has two contrasting functions in cochlear HC regeneration. On the one hand, JAG1 acts as a “mild” HC-fate repressor, and on the other hand, JAG1 is critical for preserving “progenitor-like characteristics” of cochlear SCs, a feature we previously found to be critical for cochlear HC regeneration^28–30^. Our transcriptomic and mechanistic studies reveal that JAG1 maintains the expression of pro-growth genes and pathways (e.g., PI3K-Akt-mTOR) in cochlear SCs and identify Notch1 and Notch2 as mediating JAG1’s pro-growth function. Supporting a positive role in HC regeneration, we find that after the onset of maturation (P5 and later), when cochlear SCs are already resistant to HC-fate-inducing cues, stimulation of JAG1/Notch signaling using JAG1 peptide restores the ability of SCs to generate HCs in response to HC-fate inducing cues in cochlear explants and in vivo. The positive effect of JAG1 peptide on HC-formation can be blocked by mTORC1 inhibitor rapamycin, revealing a molecular link between JAG1/Notch signaling and mTOR signaling.

## RESULTS

### Notch ligand JAG1 enhances the mitotic potential of cochlear SCs

Organoid-type cultures are well-suited to characterize the mitotic and regenerative potential of cochlear SCs^29,31^ and the SC-like Kölliker’s cells ^32^(alternative name: greater epithelial ridge cells). Kölliker’s cells (KCs) are a transient group of epithelial cells located medial to the inner HCs (IHC) and their surrounding SCs ^33^ (see schematic Fig.1a). While inner and outer HCs (IHCs, OHCs) rapidly die upon dissociation of the cochlear sensory epithelium, SCs and KCs survive dissociation, and can be propagated in 3D-extracellular matrix (Matrigel) in a cocktail of growth factors and small molecule inhibitors as spherical organoids^28^. For our experiments, we expanded organoids in culture media containing high concentrations of growth factors EGF and FGF2, valproic acid (VPA, HDAC inhibitor), TGFBR1 inhibitor (616452), and GSK-3β inhibitor (CHIR99021, activates Wnt-signaling) (Fig.1 b).

**Fig. 1.**
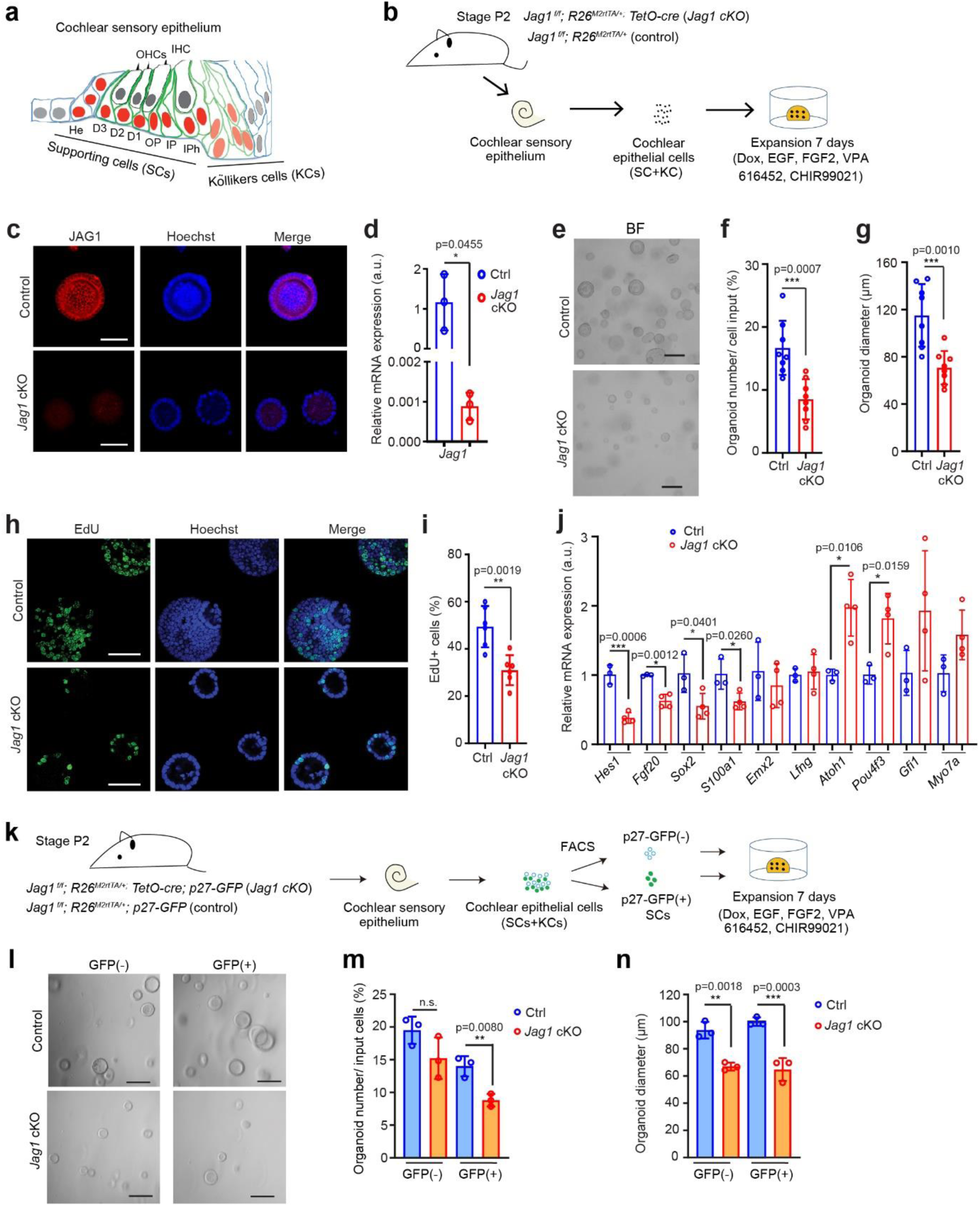
Loss of *Jag1* reduces cell cycle re-entry and proliferation of perinatal cochlear SCs and KCs in organoid culture. **a** Schematic of early postnatal murine cochlear sensory epithelium. SC-subtypes (He, Hensen cells; DC1-3 Deiter’s cells 1-3, OP, outer pillar cell; IP, inner pillar cell; iPh, inner phalangeal cells) highly express SOX2 (bright red nuclei) and JAG1 (dark green membrane). KCs express lower levels of SOX2 (faint red nuclei) and JAG1 (light green membrane). **b** Experimental scheme (**c-j**). Cochlear epithelial cells from stage P2 mice were used to establish control and *Jag1* cKO organoid cultures, which were analyzed on day 7 of expansion. **c** Confocal images of control and *Jag1* cKO organoids stained for JAG1 protein (red). Hoechst labels nuclei (blue). Scale bars=50µm. **d** RT-qPCR of relative *Jag1* mRNA expression in control (Ctrl, blue bar) and *Jag1* cKO (red bar) organoids (graphed are individual data points and mean± SD, n=3 mice per group). **e** BF images of control and *Jag1* cKO organoids. Scale bars= 200µm. **f, g** Organoid forming efficiency **(f**) and organoid diameter (**g**) for **e** (graphed are individual data points and mean ± SD, n=8 animals per group). **h** Cell proliferation in control and *Jag1* cKO organoids on day 7. Hoechst labels cell nuclei (blue) Scale bars=50µm. **i** Percentage of EdU+ cells in (**h**) (graphed are individual data points and mean ± SD, n=6 animals per group). **j** RT-qPCR of Notch target *Hes1*, SC markers (*Fgf20*, *Sox2, S100a1, Lfng*) and HC markers (*Atoh1*, *Pou4f3*, *Gfi1*, *Myo7a*) in control and *Jag1* cKO organoids (graphed are individual data points and mean ± SD, n=3 mice per group). **k** Experimental scheme (l-m). FACS-purified p27-GFP(+) SCs and p27-GFP(-) KCs from stage P2 mice were used to establish control and *Jag1* cKO organoid cultures, which were analyzed on day 7 of expansion. **l** BF images of control and *Jag1* cKO organoid cultures established from GFP(-) KCs and GFP(+) SCs. **m, n** Colony (organoid) forming efficiency (**m**) and organoid diameter (n) for e (graphed are individual data points and mean ± SD, n=3 independent cultures per group. A two-tailed, unpaired *t* test was used to calculate *P* values.

To determine whether the loss of *Jag1* affects the mitotic potential of cochlear SCs and KCs, we acutely deleted *Jag1* in cochlear organoids using a doxycycline-inducible Cre strategy. Briefly, we isolated cochlear epithelial cells (containing both SCs and KCs) from stage P2 *TetO-Cre; R26 ^rtTA*M2^*; *Jag1^f/f^* mice and littermates that lacked the *TetO-Cre* transgene (control) and cultured the cells as organoids in the presence of doxycycline (dox) (Fig.1 b). We confirmed *Jag1* deletion using RT-qPCR and anti-JAG1 immunostaining, which showed high expression of JAG1 protein in control organoids, whereas cells in *Jag1* cKO organoids lacked JAG1 protein (Fig.1 c). Similarly, RT-qPCR showed that *Jag1* transcript in *Jag1* cKO cultures was more than 100-fold less abundant than in control cultures (Fig.1 d).

After 7 days of expansion, we analyzed organoid formation efficiency (number of organoids per input cells) and organoid size (organoid diameter) in control cultures and *Jag1* cKO cultures (Fig.1 e). We found that organoid formation efficiency in control cultures was about 2-fold higher compared to *Jag1* cKO cultures (Fig.1 e and f), and the diameter of organoids in control cultures is about 1.5-fold larger compared to *Jag1* KO cultures (Fig.1 e and g). 3-hour EdU pulse experiments revealed a 1.6-fold lower rate of EdU incorporation in *Jag1* cKO organoid cultures compared to control cultures, suggesting that the smaller organoid size in *Jag1* cKO organoid cultures is due to a reduction in cell proliferation (Fig.1 h and i). RT-qPCR analysis revealed that loss of *Jag1* significantly reduced mRNA expression of *Fgf20*, *Sox2,* and *Hes1* and modestly increased *Atoh1 and Pou4f3 mRNA expression* (Fig.1 j). During cochlear development *Fgf20*, *Sox2* and *Hes1* are critical for pro-sensory cell specification (*Fgf20*, *Sox2*) ^7,34,35^ and proliferation (*Fgf20*, *Hes1*) and preventing pre-mature HC-fate induction (*Hes1*)^36^.

To determine whether loss of *Jag1* has a differential effect on SCs versus KCs, we established control and *Jag1* cKO organoid cultures with FACS-purified SCs and KCs (Fig.1k). We made use of *p27-GFP* transgene, which is highly expressed in SCs but only weakly in KCs to fractionate cochlear epithelial cells into GFP^(+)^ SCs and GFP^(-)^ KCs ^37^ (see Supplementary Fig.1 a for gating). We found that deletion of *Jag1* in KC-derived organoid cultures did not alter organoid formation (Fig. 1 l, m, GFP^(-)^) but significantly reduced average organoid size (Fig. 1 l, n, GFP^(-)^). By contrast, deletion of *Jag1* in SC-derived organoid cultures resulted in both significantly fewer (Fig.1 l, m, GFP^(+)^) and significantly smaller organoids (Fig. 1 l, n, GFP^(+)^). Taken together, our cell type-specific analysis reveals that both SCs and KCs require JAG1 for organoid growth. However, in contrast to SCs, which depend on JAG1 for organoid formation, KCs don’t require JAG1 for organoid formation. The difference in JAG1 dependency and the higher rate of organoid formation observed for KC-derived cultures is likely because in vivo KCs are still proliferating at a low rate, while cochlear SCs have permanently withdrawn from the cell cycle and are post-mitotic.

### Notch ligand JAG1 participates in cochlear HC-fate repression

Studies in mice found that inhibition of Notch signaling with GSIs in the HC-damaged cochlea stimulates trans-differentiation of immature and, to a much lesser extent, mature cochlear SCs into HCs^22,23^. To determine whether loss of *Jag1* enhances the rate of HC formation in cochlear organoids, we established control and *Jag1* cKO cochlear organoid cultures from cochlear epithelial cells and after 11 days of expansion switched to differentiation media to induce HC formation (Fig. 2a). Wnt activation (e.g., CHIR99021) in combination with Notch inhibition (e.g., LY411575) is typically used to induce HC formation in perinatal cochlear explants and organoids^28,31^. To avoid masking JAG1’s potentially HC-repressive function, LY411475 was omitted from our differentiation media (CHIR99021 only) (Fig.2a).

**Fig. 2.**
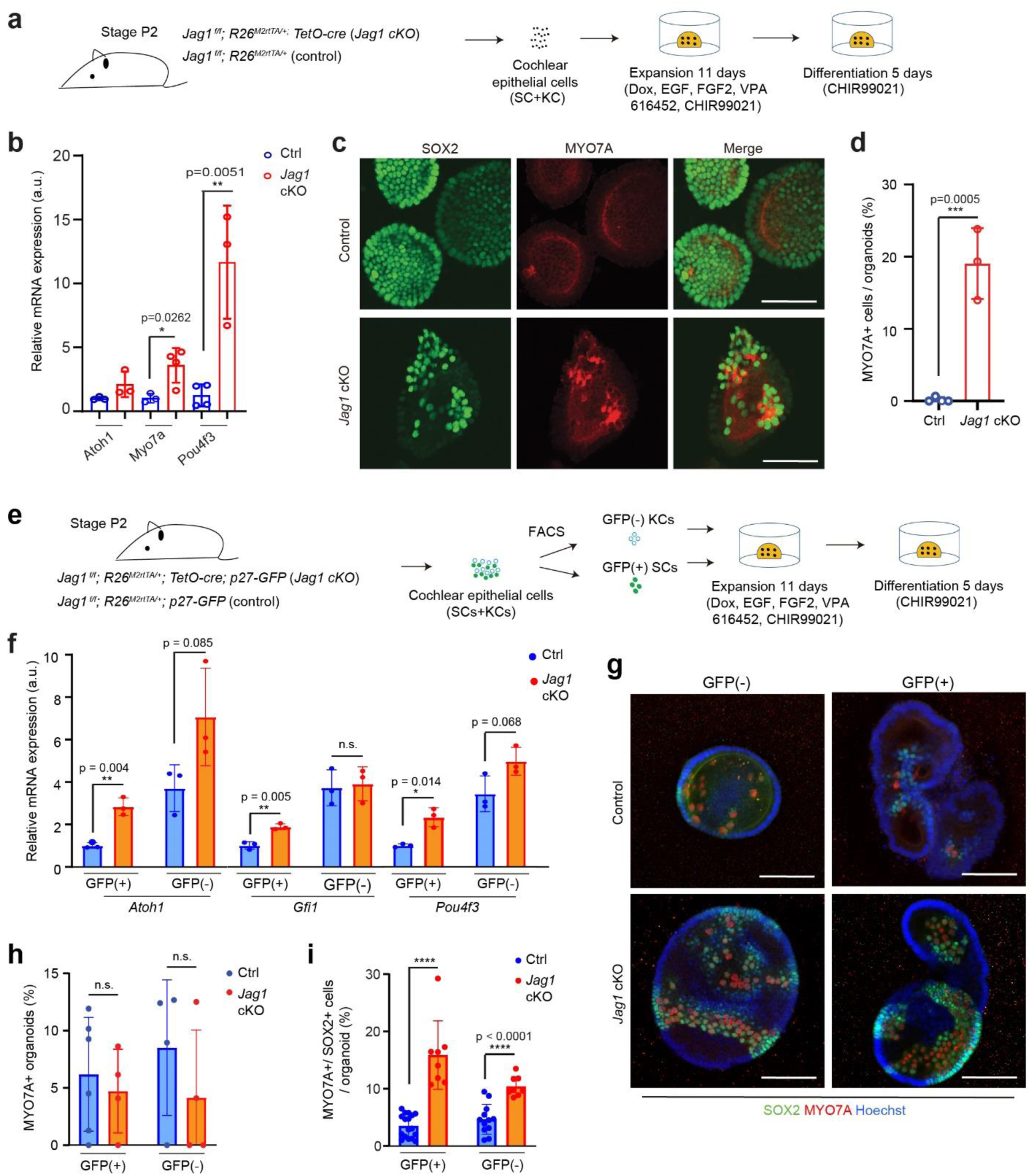
Loss of *Jag1* enhances the rate of HC formation by perinatal cochlear SCs and KCs in organoid culture. **a** Experimental scheme (b-d). Cochlear epithelial cells from stage P2 mice were used to establish control and *Jag1* cKO organoid cultures. HC formation was analyzed using RT-qPCR and immuno-staining after 5 days of differentiation. **b** RT-qPCR was used to analyze relative mRNA expression of early HC genes (*Atoh1*, *Pou4f3*, *Myo7a*) in control and *Jag1* cKO organoids (graphed are individual data points and mean ± SD, n=3 animals per group). **c** Confocal images of MYO7A (red) and SOX2 (green) stained control and *Jag1* cKO organoids. Note MYO7A and SOX2 co-staining labels new HCs. Control organoids only contain few scattered new HCs whereas *Jag1* cKO organoids contain clusters of new HCs. Scale bars=50µm. **d** Quantification of percentage of HCs per control and *Jag1* cKO organoid in (c). **Experimental scheme** (f-i) FACS-purified P27-GFP^(+)^ SCs and p27-GFP^(-)^ KCs from stage P2 mice were used to establish control and *Jag1* cKO organoid cultures. HC formation was analyzed by RT-qPCR and immuno-staining after 5 days of differentiation. **f** RT-qPCR was used to analyze relative mRNA expression of early HC-specific transcription factors (*Atoh1*, *Gfi1*, *Pou4f3*) in control and *Jag1* cKO organoids (graphed are individual data points and mean ± SD, n=3 independent biological replicates per group). **g** Confocal images of KC-derived (GFP(-)) and SC-derived (GFP(+)) control and *Jag1* cKO organoids stained for MYO7A (red) and SOX2 (green) Scale bars=100µm. **h, i** Percentage of HC-containing organoids (**h**) and percentage of HCs in HC-containing organoids (**i**) in control and *Jag1* cKO organoid cultures (graphed are individual data points and mean ± SD, n=3 independent biological replicates per group). A two-tailed, unpaired *t* test was used to calculate *P* values. \**P* ≤ 0.05, \*\**P* <0.01, \*\*\**P* <0.001.

Consistent with JAG1 participating in HC-fate repression, we found that at day 5 of differentiation, *Jag1* cKO organoids expressed the transcripts of HC-specific *Myo7a* gene at more than 3–fold higher level and HC-specific *Pou4f3* gene at a more than 10-fold higher level than control organoids (Fig.2 b), indicating a higher rate of HC production in *Jag1* cKO organoid cultures compared to control cultures. Next, we used immunostaining against MYO7A and SOX2 to visualize newly formed HCs in organoids. Nascent cochlear HCs co-express MYO7A and SOX2, distinguishing them from existing HCs, which lack SOX2 expression^38^. We found that in *Jag1* cKO cultures, organoids contained large clusters of MYO7A^+^SOX2^+^ HCs, with HCs constituting 20% of total cells per organoid, whereas, in control cultures, organoids contained only a few scattered HCs (Fig. 2 c and d).

To determine whether *Jag1* deficiency differentially affects HC formation by SCs and KCs, we established control and *Jag1* cKO organoids with p27-GFP^(+)^ SCs and p27-GFP^(-)^ KCs (Fig.2e), and after 5 days of differentiation, analyzed HC formation using RT-qPCR and immunostaining. We found that in the absence of *Jag1,* SC-derived cultures expressed early HC-specific transcripts (*Atoh1*, *Pou4f3*, *Gfi1*) 2-3-fold higher than corresponding control cultures (Fig.2 f, GFP^(+)^). By contrast, deletion of *Jag1* in KC-derived cultures resulted only in a modest increase of *Atoh1* and *Pou4f3* expression (Fig.2 f, GFP^(-)^). MYO7A and SOX2 immunostaining revealed that the percentage of organoids that contained HCs was similar across the four conditions (Fig.2 h). However, as anticipated, the percentage of HCs per organoid was significantly higher in both SC and KC-derived *Jag1* cKO organoid cultures compared to control cultures, and only *Jag1* cKO organoids contained small to mid-size clusters of HCs (Fig.2 g, i).

To further investigate the role of JAG1 in HC-fate repression, we analyzed whether loss of *Jag1* increases the rate of HC regeneration in early postnatal cochlear explants. We harvested cochleae from stage P2 *TetO-Cre*; *Jag1^f/f^; R26^M2-rtTA/+^*mice and *Jag1^f/f^; R26^M2-rtTA/+^* control littermates that received dox starting at E18.5 and used microdissection to isolate the cochlear sensory epithelium and underlying mesenchyme and spiral ganglion (Supplementary Fig.2a), To ablate HCs in control and *Jag1* cKO cochlear explants gentamicin was added for the first 15 hours of culture. We used a dose of 100 µg/mL gentamicin, which is highly effective in killing cochlear HCs throughout the mid-apical to basal cochlea without damaging surrounding SCs^22^. After 15 hours, gentamicin-containing culture media was removed and *Jag1* cKO and control explants were cultured with CHIR99021 or DMSO (0.05%; vehicle control) for 3 more days (Supplementary Fig.2b). On day 4 (∼P6) explants were stained for SOX2 and MYO7A and EdU incorporation and new HC formation (MYO7A^+^SOX2^+^) was analyzed in the mid-apex. We found that the rate of “spontaneous” HC regeneration was extremely low in both DMSO-treated *Jag1* cKO and control explants (4-6 HCs/ 200 µm = ∼4%-6% of total HCs in undamaged cochlea) (Supplementary Fig.2c and d). Moreover, the number of SCs that incorporated EdU was extremely low in DMSO-treated control and *Jag1* cKO explants (Supplementary Fig.2c, e), and none of the newly formed HCs incorporated EdU, indicating that the few spontaneously regenerated HCs were generated through non-mitotic mechanisms (Supplementary Fig.2c, f). The presence of CHIR99021 (CHIR) increased the rate of cochlear SC proliferation in both control and *Jag1* cKO explants to a similar extent (∼30 SOX2^+^ EdU^+^ cells/ 200 µm = ∼16% of total SCs) (Supplementary Fig.2c, e). Furthermore, in the presence of CHIR99021 modestly increased HC-production in *Jag1* cKO explants (∼8 HC/ 200 µm = ∼8% of total HCs in undamaged cochlea) (Supplementary Fig.2c, d) and about half of the newly formed HCs in *Jag1* cKO cultures derived from dividing cells (Supplementary Fig.2c, f). Our data, taken together, indicates that JAG1/Notch signaling participates in HC-fate repression. However, the effect of *Jag1* deletion on new HC-formation is modest, hinting at a more complex role in this process.

### Notch ligand JAG1 maintains metabolic and progenitor genes in cochlear SCs and KCs

To identify genes and pathways that are regulated by JAG1/Notch signaling in cochlear organoids, we established control and *Jag1* cKO organoids from P2 cochlear epithelial cells and analyzed their transcriptome after 7 days of expansion using RNA sequencing (RNA_seq) (Fig.3a) (Supplementary Table 1). We first examined Notch ligands and receptors’ expression in control organoids. We found that *Jag1, Notch1*, *Notch2,* and *Notch3* transcripts were highly expressed. Consistent with organoids being close to void of HCs, HC-specific ligand *Dll1* transcript was close to undetectable, and HC-specific ligand *Jag2* transcript was more than 100-fold lower expressed than *Jag1* (Supplementary Fig.3a). Next, we used the data to identify differentially expressed genes (DEG). Our analysis identified 802 DEGs (q-value 0.01), with 160 genes upregulated and 642 genes downregulated in *Jag1* CKO organoids compared to control organoids (Supplementary Table S2). As anticipated, transcripts for *Jag1* and down-stream Notch pathway effector genes (*Hes1*, *Hey1, HeyL, Sox2*) were among the top down-regulated transcripts in *Jag1* cKO organoid cultures compared to control organoid cultures (Fig. 3b, d) (Supplementary Fig.3b). In addition, transcripts of growth-promoting genes (e.g., *Fgf10*, *Igf2, Igfbp3, Ntf3*), and genes regulating cell metabolism and metabolite transport (e.g., *Me2*, *Eif4ebp1*, *Slc16a3*, *Slc7a5*) were significantly downregulated in *Jag1* CKO organoids compared to control organoids (Fig.3b) (Supplementary Table 2). Among the cohort of genes that were significantly upregulated in *Jag1* CKO organoids compared to control were genes expressed in HCs (e.g. *Tunar*^39^, *Pou4f3, Scx*^40^) and genes expressed in interdental cells (e.g. *Smoc2*^39^ and *Otoa*^41^), which are a group of cochlear epithelial cells critical for the production of tectorial membrane proteins (Fig. 3b)(Supplementary Table 2). To identify pathways and biological processes that may be disrupted by the loss of JAG1-mediated signaling, we performed gene ontology enrichment analysis on the list of down-regulated genes using Metascape, a web-based portal^42^. As anticipated ‘Notch signaling pathway’ and ‘regulation of nervous system development’ were among the top 20 pathways and biological processes (Fig.3c) (Supplementary Table 3). However, 5 out of the 10 top scoring pathways and biological processes were related to cell stress response (response to endoplasmic reticulum stress, HIF-1 pathway) and cell metabolism (PI3K-Akt-signaling pathway, metabolism of carbohydrates, amino acid metabolic process) (Supplementary Fig. 4a-c) (Fig.3c) (Supplementary Table S3), suggesting that loss of *Jag1* disrupted PI3K-Akt-mTOR signaling in cochlear SCs and KCs.

**Fig. 3.**
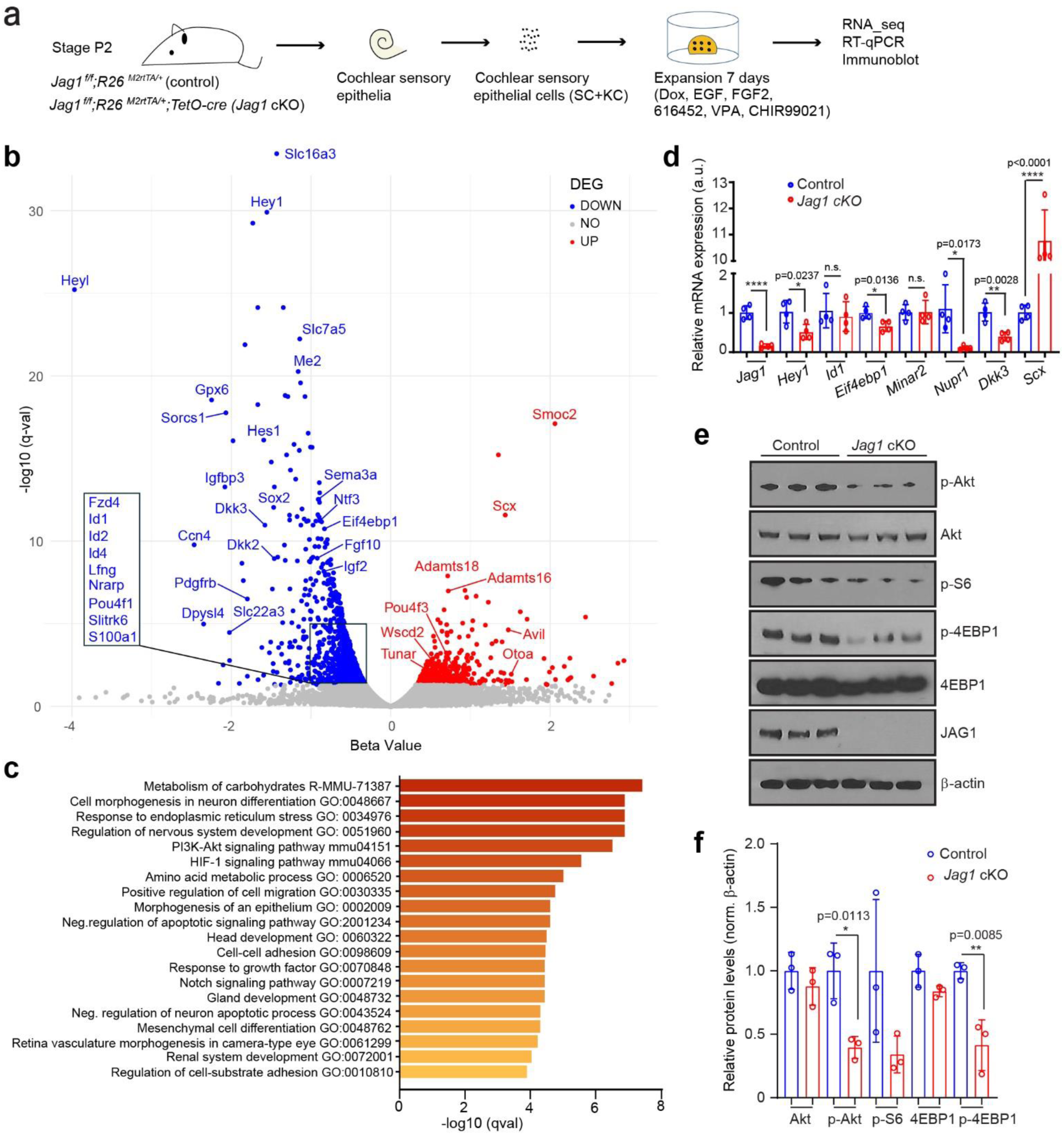
**Loss of *Jag1* reduces the expression of progenitor and SC-specific genes and attenuates PI3K-Akt-mTOR signaling**. **a** Experimental scheme. Cochlear epithelial cells from stage P2 mice were used to establish control and *Jag1* cKO organoid cultures, which were analyzed on day 7 of expansion. **b** Volcano plot of RNA-seq data. Plotted is beta-value (x-axis) versus −log10 q-value (y-axis). Transcripts that are significantly up regulated in response to *Jag1* cKO are marked in red dots, and transcripts that are significantly downregulated are marked in blue dots. **c** Biological processes and pathways associated with *Jag1* cKO downregulated genes ranked by adjusted *P* value (*q* value). **d** RT-qPCR was used to analyze relative mRNA expression of *Jag1*, *Hey1*, *Id1*, *Eif4ebp1*, *Minar2*, *Nupr1*, *Dkk3*, *Scx* in control and *Jag1* cKO organoids (graphed are individual data points and mean ± SD, n=4 animals). **e** Immunoblots of p-Akt, Akt, p-S6, p-4EBP1, 4EBP1, JAG1 and β-actin proteins in control and *Jag1* cKO organoids. **f** Normalized p-Akt, p-S6 and p-4EBP1 protein levels in (**e**) (graphed are individual data points and mean ± SD, n = 3 animals, from one representative experiment). Two-tailed, unpaired Student’s *t* test was used to calculate *P* values. \**P* ≤ 0.05, \*\**P* <0.01, \*\*\**P* <0.001 and \*\*\*\**P* < 0.0001.

### Notch ligand JAG1 maintains PI3K-Akt-mTOR signaling in cochlear SCs and KCs

To determine whether PI3K-Akt-mTOR signaling is downregulated in *Jag1* cKO organoids, we analyze the expression levels of phosphorylated (p) forms of kinase Akt (p-Akt, Ser473) ribosomal protein S6 (p-S6, Ser240/244) and eukaryotic initiation factor 4E binding protein 1 (p-4EBP1, Thr37/46) proteins in control and *Jag1* cKO organoid protein extracts by immunoblot. An increase in the protein levels of the p-S6 (Ser240/244) and p-4EBP1 (Thr37/46)^43^ correlate with increased mTOR activity, whereas p-Akt (Ser473) is a readout for maximal Akt activation downstream of PI3K^44^. Our analysis revealed that protein levels of p-AKT and p-4EBP1 were significantly reduced in *Jag1* cKO organoids compared to control, identifying JAG1 as a positive regulator of PI3K-AKT-mTOR signaling (Fig.3 e and f).

JAG1 peptides that contain the extracellular domain of human JAG1 fused in frame with Fc sequence (JAG1-Fc) have been shown to efficiently activate Notch signaling in various contexts^45–47^. To determine whether extracellular JAG1 can rescue the defects in organoid formation and growth observed in *Jag1* CKO organoids, we established P2 *Jag1* cKO and control organoids and cultured them with (5, 50, 500 ng/mL) or without (0 ng/mL) JAG1-Fc peptide (Ser32--Ser1046)^48^(Fig. 4a). We found that the rate of organoid formation (Fig.4 b, c) and organoid growth (Fig. 4 b, d) was significantly increased by exposure to JAG1-Fc peptide in both control and *Jag1* cKO organoid cultures. However, likely due to endogenous JAG1 ligand competing for Notch receptor binding, a higher concentration of JAG1-Fc peptide was required to solicit a pro-growth response in control organoids (50 ng/mL) compared to *Jag1* cKO organoids (5 ng/mL) (Fig.4, b-d). Correlating with the higher rate of organoid formation and organoid growth, the JAG1-Fc peptide increased the abundance of p-Akt and p-4EBP1 protein in both control and *Jag1* CKO organoids (Fig.4 e). The transcription of *Jag1* in cochlear SCs^25^ is positively regulated by Notch signaling^49^. Consistent with JAG1-Fc activating Notch receptor signaling, exposure to JAG1-Fc peptide led to a dose-dependent increase in endogenous JAG1 protein expression in control organoids (Fig.4 e, control).

**Fig. 4.**
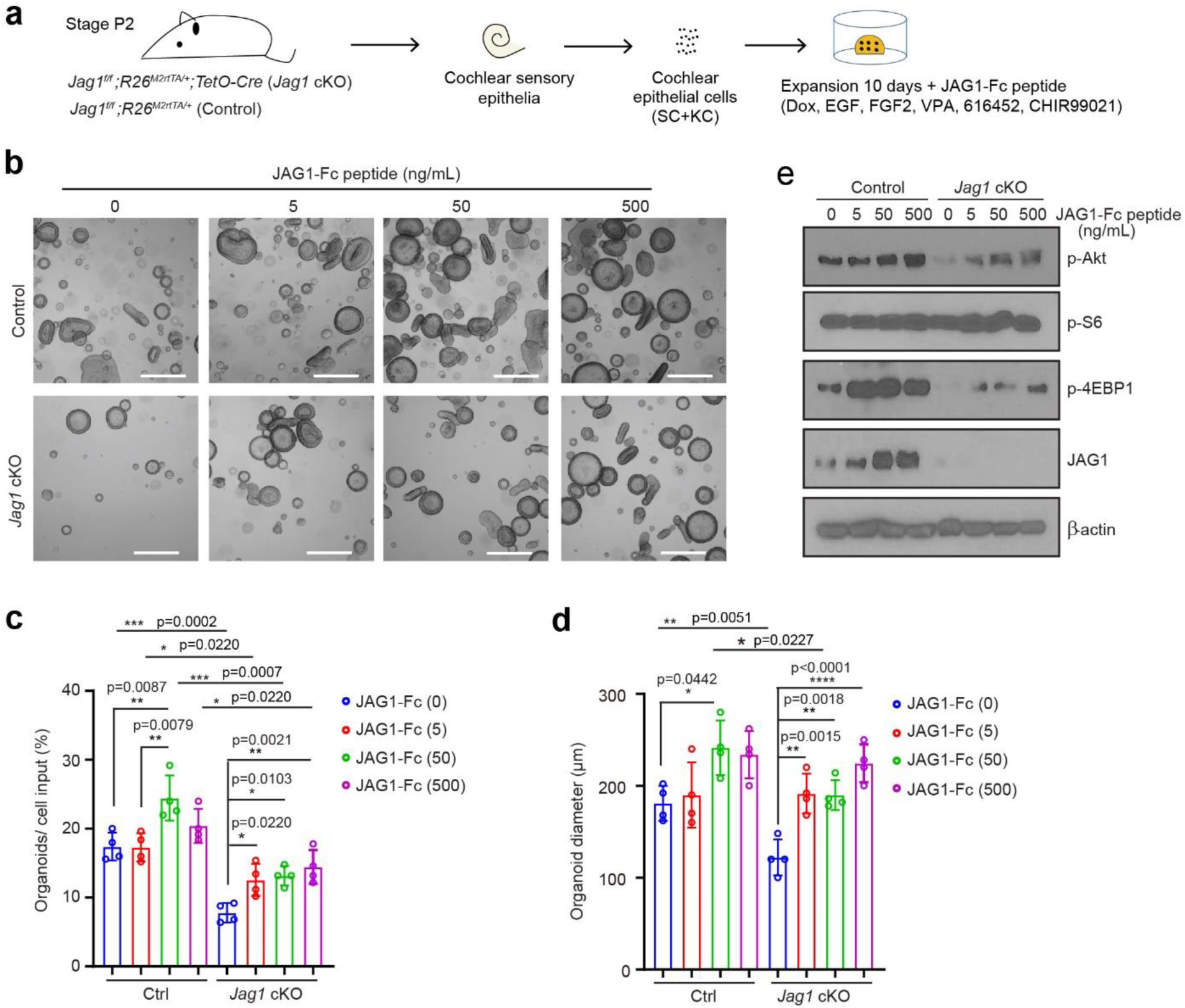
Exogenous JAG1-peptide restores growth and PI3K-Akt-mTOR signaling to near normal levels in *Jag1* deficient cochlear organoids. **a** Experimental scheme. Cochlear epithelial cells from stage P2 mice were used to establish control and *Jag1* cKO organoid cultures, which were analyzed on day 10 of expansion. Organoids were treated with JAG1-Fc peptide on day 3 of expansion and cultured for 7 additional days. **b** BF images of control and *Jag1* cKO organoids treated with or without JAG1-Fc peptide on day 10 of expansion. Scale bars=400µm. **c-d** Quantification of organoid forming efficiency (**c**) and organoid diameters (**d**) for **b** (graphed are individual data points and mean ± SD, n=4 animals per group). **e** Immunoblot of p-Akt, pS6, p-4EBP1, JAG1 and β-actin protein in control and JAG1 peptide treated organoids on day 10 of expansion. One-way ANOVA with Tukey’s correction was used to calculate *P* values. \**P* ≤ 0.05, \*\**P* <0.01 and \*\*\*\**P* < 0.0001.

To independently confirm that stimulating JAG1-Notch signaling promotes organoid formation and organoid growth, we overexpressed full-length murine JAG1 protein in P2 cochlear organoids using a lentiviral strategy. To mark infected cells, control, and JAG1 expressing lentivirus expressed red-fluorescent protein mCherry (Supplementary Fig. 5a). Our analysis revealed that JAG1 overexpression significantly increased organoid size and organoid formation efficiency (Supplementary Fig.5b-d) and increased the rate of cell proliferation (EdU^+^ mCherry^+^ cells/ organoid) (Supplementary Fig.5e and f). In summary, our data indicates that JAG1 positively regulates the mitotic capacity of early postnatal cochlear SCs and KCs.

Qualitative similar positive effects on cochlear organoid growth and organoid formation were observed with low activation of Notch1 receptor signaling. To stimulate Notch1 receptor signaling, we infected P2 wild-type cochlear epithelial cells with lentivirus that expressed N1ICD in a dox-dependent manner and expanded organoids with no dox (0), low (0.25, 0.5) or high dox (5, 10 μg/mL) for 7 days (Fig. 5a). We found that relative low concentration of dox (0.5 μg/mL) significantly increased organoid size (Fig. 5 b, c) and organoid formation efficiency (Fig. 5 b, d) compared to no dox (0 μg/mL) while 10-20x higher concentrations of dox (5 and 10 μg/mL) had an adverse effect on organoid growth and formation, significantly decreasing organoid size and organoid formation efficiency (Fig. 5 b-d).

**Fig. 5.**
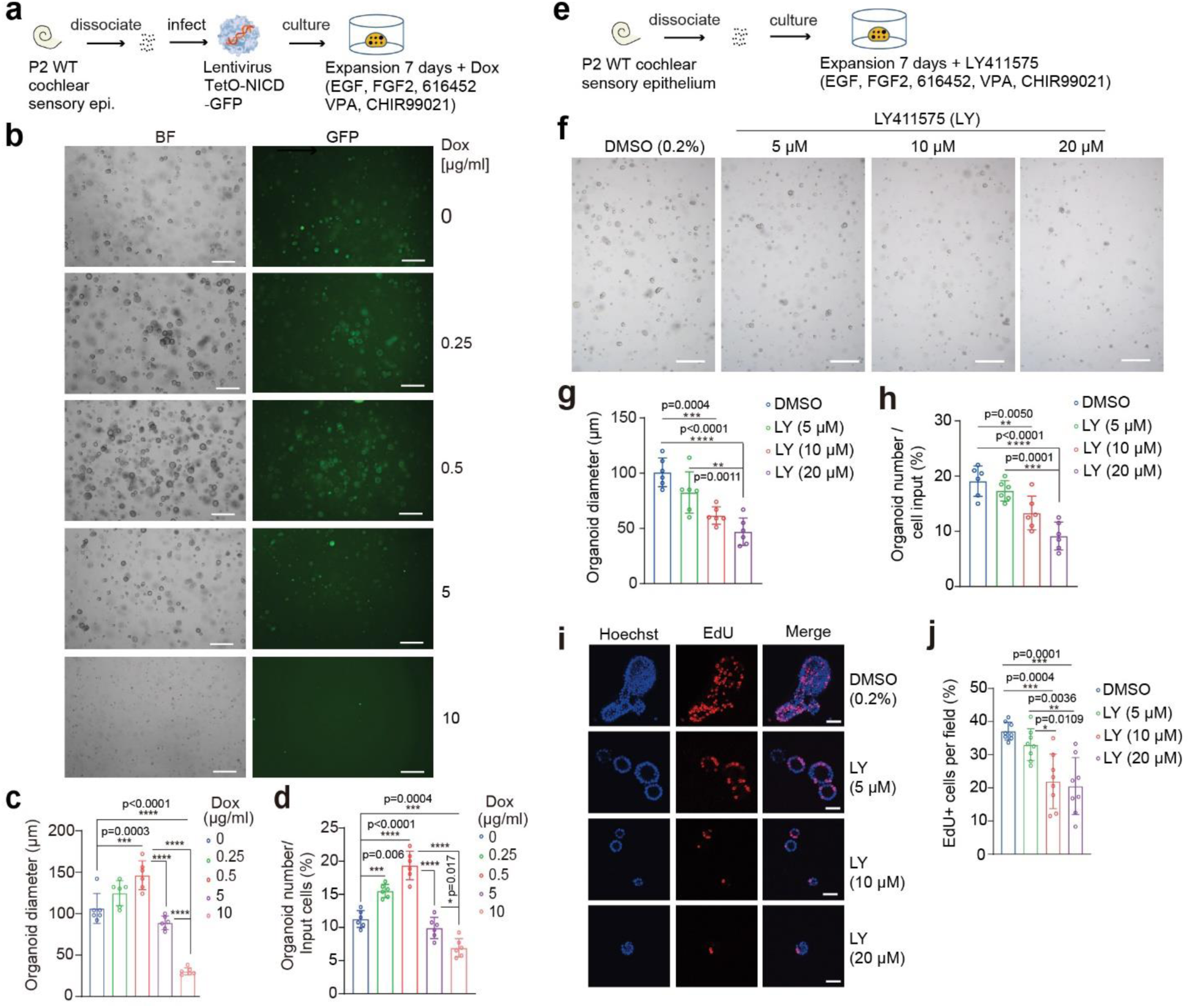
Low Notch signaling promotes cochlear SC and KC proliferation, organoid formation, and organoid growth. **a** Experimental scheme (**b-d**). Cochlear epithelial cells from stage P2 wild type mice were infected with lentivirus that expressed the intracellular domain of Notch1 (NICD) in a dox-dependent manner. Dox (0.25, 0.5, 5 and 10 µg/mL) was added to the culture medium on day 2 and organoids were analyzed after 7 days of expansion. **b** BF and GFP images of control (0 dox) and NICD expressing organoids. GFP labels organoids infected with lentivirus. Scale bars=400µm. **c-d** Organoid diameter (**c**) and organoid forming efficiency (**d**) for **b** (graphed are individual data points and mean ± SD, n=6 independent biological replicates, two independent experiments). **e** Experimental scheme (**f-g**). **f** BF images of DMSO (vehicle control) and LY411575 treated organoids on day 7 of expansion. LY411575 or DMSO (0.2%) was added to the culture medium on day 2 of expansion. Scale bars=400µm. **g-h** Organoid diameter (**g**) and organoid forming efficiency (**h**) for **f** (graphed are individual data points and mean ± SD n=6, two independent experiments). **i** Cell proliferation in control (DMSO) and LY411575 treated organoids. A single EdU pulse was given on day 7 of expansion and EdU incorporation (red) was analyzed 1 hour later. Hoechst labels cell nuclei (blue). Scale bars=25µm. **j** Percentage of EdU+ cells in (**i**) (graphed are individual data points and mean ± SD, n=8 animals, two independent experiments). One-way ANOVA with Tukey’s correction was used to calculate P values. \**P* ≤ 0.05, \*\**P* <0.01, \*\*\**P*<0.001 and \*\*\*\**P* < 0.0001.

Next, we examined whether inhibition of Notch receptor signaling with LY411575 (GSI) reduces organoid formation and organoid growth. We established cochlear organoids from P2 wild-type cochlear epithelial cells and expanded organoids in the presence of DMSO (0.2%, vehicle control) or in the presence of increasing concentrations of LY411575 (5, 10, 20 μM) (Fig.5 e) for 4 days. We found that 5 μM LY411575, a dosage commonly used to induce SC-to-HC conversion in perinatal cochlear organoids or explants, did not affect organoid formation or growth, but 2 and 4-fold higher concentrations of LY411575 (10, 20 μM) significantly decreases the average organoid size (Fig. 5 f, g) and organoid formation in a dose-dependent manner (Fig. 5 f, h). EdU pulse experiments showed that cells in organoids cultured with 10 or 20 μM LY411575 incorporated EdU at a significantly lower rate than control organoids (DMSO) or organoids cultured with 5 μM LY411575 (Fig.5 i, j). In sum, these data reveal that low levels of Notch1 receptor signaling promote cochlear organoid formation and growth, while high levels of Notch1 receptor signaling are inhibitory. Furthermore, our data indicates that 5 μM of LY411575, a concentration used to induce SC-to-HC conversion, does not affect cochlear organoid formation and growth.

### Loss of *Notch1/2* reduces PI3K-Akt-mTOR signaling in cochlear SCs and KCs

Early postnatal cochlear SCs and KCs cells co-express the Notch receptors Notch1, Notch2, and Notch3^39^, suggesting that one or more of these three Notch receptors mediate the positive effects of JAG1 on organoid formation, growth, and PI3K-Akt-mTOR signaling. To identify the Notch receptor(s) through which JAG1 signals, we established P2 cochlear organoid cultures with *Notch1*, *Notch2,* or *Notch3* deficient cochlear epithelial cells (SCs + KCs) and corresponding control cochlear epithelial cells. To analyze the *Notch3* function, we used *Notch3* heterozygous (*Notch3^+/-^*) (control) and *Notch3* homozygous mutant (*Notch3^-/-^*) mice (Supplementary Fig.6 a)^50^. Analysis of cochlear HC phenotype in early postnatal *Notch3*^-/-^ mice and *Notch3^-/+^* littermates (control) revealed no developmental defects. Both control and *Notch3* deficient cochlear sensory epithelium contained a single row of inner HCs and three rows of outer HCs, surrounded by SCs (Supplementary Fig.6 b), indicating that Notch3 function *in vivo* is not required for pro-sensory cell maintenance, nor is Notch3 required for HC fate repression. We next established cochlear organoid cultures with stage P2 *Notch3^+/-^*and *Notch3^+/-^* cochlear epithelial cells and expanded them for 7 days (Supplementary Fig.6 a). Our analysis of colony-forming efficiency (organoid formation) and organoid size (diameter) revealed no differences between *Notch3^-/+^*and *Notch3^-/-^* organoid cultures (Supplementary Fig.6 c-e). RT-qPCR experiments revealed that *Notch3^-/-^* organoids expressed Notch target genes *Hes1*, *Hes5,* and *Sox2* at lower levels than *Notch3^-/+^* (control organoids), however *Atoh1* expression remained unchanged (Supplementary Fig.6 f). Furthermore, immunoblots showed that p-Akt, p-S6, and p-4EBP1 protein expression in *Notch3^-/-^* organoids were unchanged compared to *Notch3^-/+^* organoids (Supplementary Fig.6 g and h).

Previous studies have shown that Notch1 is critical for HC fate repression, but the role of Notch1 signaling in cochlear SC and KC proliferation and whether Notch1 is required for PI3K-Akt-mTOR signaling is unclear. To analyze Notch1 function, we established cochlear organoids using cochlear epithelial cells from stage P2 *Sox2^CreER/+^; Notch1^f/f^* mice and *Notch ^f/f^* littermates (control). In the early postnatal cochlea, tamoxifen-inducible *Sox2^CreER^* transgene is highly expressed in SCs and KCs ^51^. To induce *Notch1* deletion in culture, 4-hydroxy-tamoxifen (4-OH TM) was included in the culture media. We also established additional control cultures with DMSO (0.02%, vehicle control) to control *Sox2* haploinsufficiency^51^ (Supplementary Fig. 7 a). Our analysis revealed that organoid formation and growth in *Notch1* cKO organoid cultures (*Sox2^CreER/+^; Notch1^f/f^* + 4-OH TM) was not significantly different compared to control cultures (*Sox2^CreER/+^; Notch1^f/f^* + DMSO or *Notch1^f/f^* + DMSO or *Notch1^f/f^* + 4-OH TM) (Supplementary Fig.7 b-d). Furthermore, analysis of p-S6, p-Akt, and p-4EBP1 protein expression in *Notch1* cKO and control organoids revealed no significant differences (Supplementary Fig.7 e and f).

Next, we analyzed the function of the Notch2 receptor using a cochlear-specific Cre line (*Emx2^Cre/+^*)^52^ to conditionally delete *Notch2* by itself or in combination with *Notch1*. Analysis of the cochlear HC phenotype of stage P4 *Notch2* deficient mice revealed no obvious HC patterning defects. Like control cochlear tissue (Cre negative), the cochlear sensory epithelia that lacked both *Notch2* alleles (*Emx2^Cre/+^; Notch2^f/f^; Notch1^fl/+^*) contained a single row of inner HCs and three rows of outer HCs (Supplementary Fig.8 a), indicating that Notch2 function in the developing cochlea is not required for the maintenance of pro-sensory cells, nor is Notch2 required for HC fate repression. By contrast, cochlear sensory epithelia that lacked both *Notch1* alleles (*Emx2^Cre/+^; Notch2^f/+^; Notch1^f/f^*) showed severe HC patterning defects, containing many newly formed HCs (MYO7A^+^SOX2^+^), which is consistent with Notch1’s known role in HC-fate repression (Supplementary Fig.8 a).

To determine whether loss of *Notch2* alters the mitotic capacity of cochlear SCs and KCs, we established cochlear organoid cultures with cochlear epithelial cells isolated from stage P2 *Emx2^Cre/+^; Notch2^f/f^* mice and control litter mates and analyzed organoid growth and formation after 10 days of expansion (Supplementary Fig.8 b). Our analysis revealed that both organoid formation efficiency and organoid size were significantly reduced in *Notch2* deficient (*Emx2^Cre/+^; Notch2^f/f^)* organoid cultures compared to control (*Notch2*^f/f^) (Supplementary Fig.8 c-e). However, immunoblots analyzing p-Akt, p-S6, and p-4EBP1 protein expression revealed no reduction in PI3K-Akt-mTOR signaling in Notch2 deficient organoids (Supplementary Fig.8 f and g) or acutely isolated Notch2 deficient cochlear sensory epithelia compared to control (Supplementary Fig. 8 h and i).

To determine whether Notch1 may have compensated for the loss of Notch2, we made use of *Notch1^fl/fl^; Notch2^fl/fl^* mice to acutely knockout both *Notch1* and *Notch2* (*Notch1/2* DKO) in cochlear organoids. We isolated cochlear epithelial cells from stage P2 *Notch1^fl/fl^; Notch2^fl/f^* mice and infected the cells with control lentivirus or Cre expressing lentivirus, after which we expanded them as organoids for 10 days (Fig.6 a). To track transduced cells, both control and Cre expressing lentivirus expressed the red fluorescent protein mCherry (Fig. 6 b). RT-qPCR revealed that Cre-infected organoid cultures (*Notch1/2* DKO) expressed *Notch1*, *Notch2*, and Notch effector genes *Hes1*, *Hey1,* and *HeyL* at a 2-3-fold lower level than organoid cultures infected with control virus (Control), confirming successful disruption of Notch1/2 signaling (Fig. 6 c). Furthermore, we found that *Notch1/2* DKO cultures formed 2-fold fewer organoids than control cultures (Fig.6 b and d), and the diameter of organoids in *Notch1/2* DKO cultures was about 1.5-fold smaller compared to organoids in control cultures (Fig. 6 b and e). Furthermore, analysis of p-S6, p-Akt and p-4EBP1 protein expression using immunoblots revealed that the protein levels of p-Akt and p-4EBP1 were significantly lower in *Notch1/2* DKO organoids than control organoids (Fig. 6 f and g) indicating that Notch1 and Notch2 have redundant functions in regulate PI3K-Akt-mTOR signaling in cochlear SCs and KCs.

**Fig. 6.**
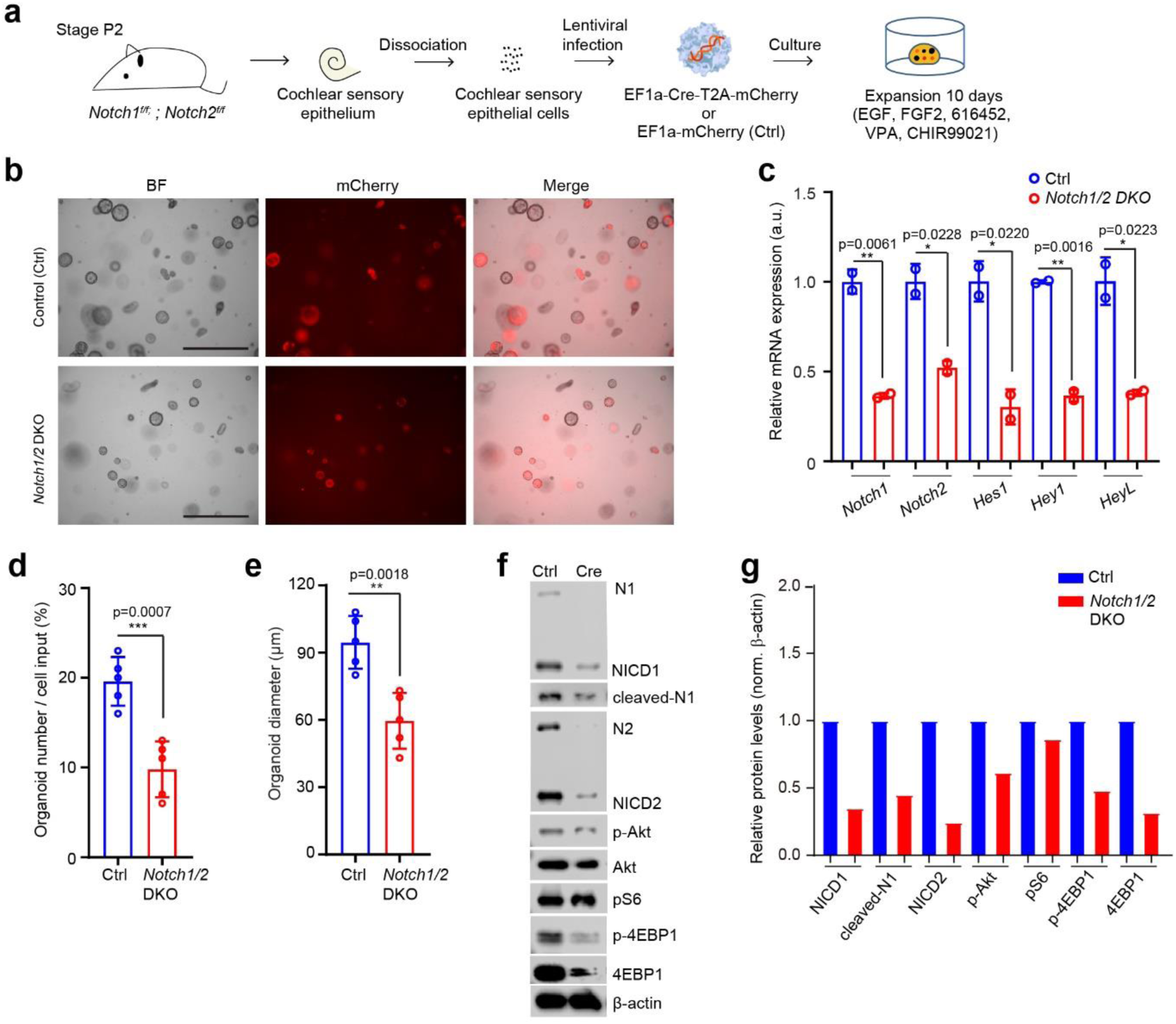
Loss of *Notch1* and *Notch2* reduces formation and growth of cochlear organoids and attenuates PI3K-Akt-mTOR signaling. **a** Experimental scheme. Organoid cultures were established with cochlear epithelial cells from stage P2 *Notch1^f/f^ Notch2^f/f^* mice. Cells were infected with Cre–mCherry and mCherry only (control) expressing lentivirus (LV) prior plating and organoids expanded for 10 days. **b** BF images of control and *Notch1/2* double knockout (DKO) organoids analyzed on day of expansion. Scale bars=400µm. **c** RT-qPCR analyzing relative mRNA expression of *Notch1, Notch2* and Notch effectors (*Hes1*, *Hey1* and *HeyL*) in control and *Notch1/2* DKO organoids on day 10 of expansion (graphed are individual data points and mean ± SD, n=3 independent biological replicates, two independent experiments). **d-e** Organoid (colony) forming efficiency (**d**) and organoid diameter (**e**) was analyzed for experiment shown in **b** (graphed are individual data points and mean ± SD, n=5 independent biological replicates, two independent experiments). **f** Immunoblot of Notch1 (N1), NICD1, cleaved N1, Notch2 (N2), NICD2, p-Akt, Akt, p-S6, p-4EBP1, 4EBP1 and β-actin proteins in control and *Notch1/2* dKO cochlear organoids on day 10 of expansion. **g** Bar graph showing normalized protein levels of Notch1 (N1), NICD1, cleaved N1, Notch2 (N2), NICD2, p-Akt, Akt, p-S6, p-4EBP1, 4EBP1 in (**f**). Two-tailed, unpaired Student’s *t* test was used to calculate *P* values. \**P* ≤ 0.05, \*\**P* <0.01 and \*\*\**P* < 0.001.

To further confirm the growth-inhibitory effects of *Notch1* and/or *Notch2* deletion, we knocked down *Notch1* and/or *Notch2* expression in SC-derived organoids using short hairpin RNAs (shRNAs) (Supplementary Fig.9 a). We designed three shRNA constructs each for targeting *Notch1* or *Notch2* and used RT-qPCR to analyze their knockdown efficiency in pilot experiments. Based on these experiments, we selected *shNotch1-2* and *shNotch2-3* to knock down *Notch1* and/or *Notch2* in subsequent experiments (Supplementary Fig. 9 b, c). We isolated cochlear SCs from stage P2 *Lfng-GFP* transgenic mice using FACS as previously described ^29^ (see Supplementary Fig.1 b for gating). Following lentiviral infection, Lfng-GFP^(+)^ cochlear SCs were expanded as organoids for 7 days (Supplementary Fig. 9 d). Consistent with our *Notch1* and or *Notch2* knockout experiments that used unfractionated cochlear epithelial cells to establish organoids, we found that *Notch2* knockdown and, to a larger extent, combined *Notch1* and *Notch2* knockdown (*Notch1/2)* significantly reduced organoid formation efficiency in cochlear SC-derived organoids (Supplementary Fig. 9 d and e). Furthermore, we found that *Notch2* knockdown and *Notch1/2* knockdown significantly reduced the organoid size of cochlear SC-derived organoids (Supplementary Fig. 9 d and f). By contrast, *Notch1* knockdown did not significantly reduce organoid formation, nor did it reduce organoid size compared to control (Supplementary Fig. 9 d-f).

### Exogenous JAG1 enhances the mitotic and HC-forming potential of cochlear SCs

At the onset of cochlear maturation at around P5/P6, the ability of SCs to form HCs sharply declines, which is accompanied by a reduction in responsiveness to Notch inhibition and an overall reduction in Notch1 signaling strength ^5,24^. To characterize potential changes in *Jag1* expression during cochlear maturation, we isolated SCs from stage P1, P5, and P13 Lfng-GFP transgenic mice using FACS (see Supplementary Fig.1b for gating) and analyzed *Jag1* mRNA abundance using RT-qPCR (Fig.7 a). Our analysis revealed that *Jag1* mRNA expression in Lfng-GFP^(+)^ cochlear SCs declines between stages P1 and P5 by 2-fold and more than 5-fold between stages P1 and P13, which is around the time when mice start to hear (Fig. 7 b).

**Fig. 7.**
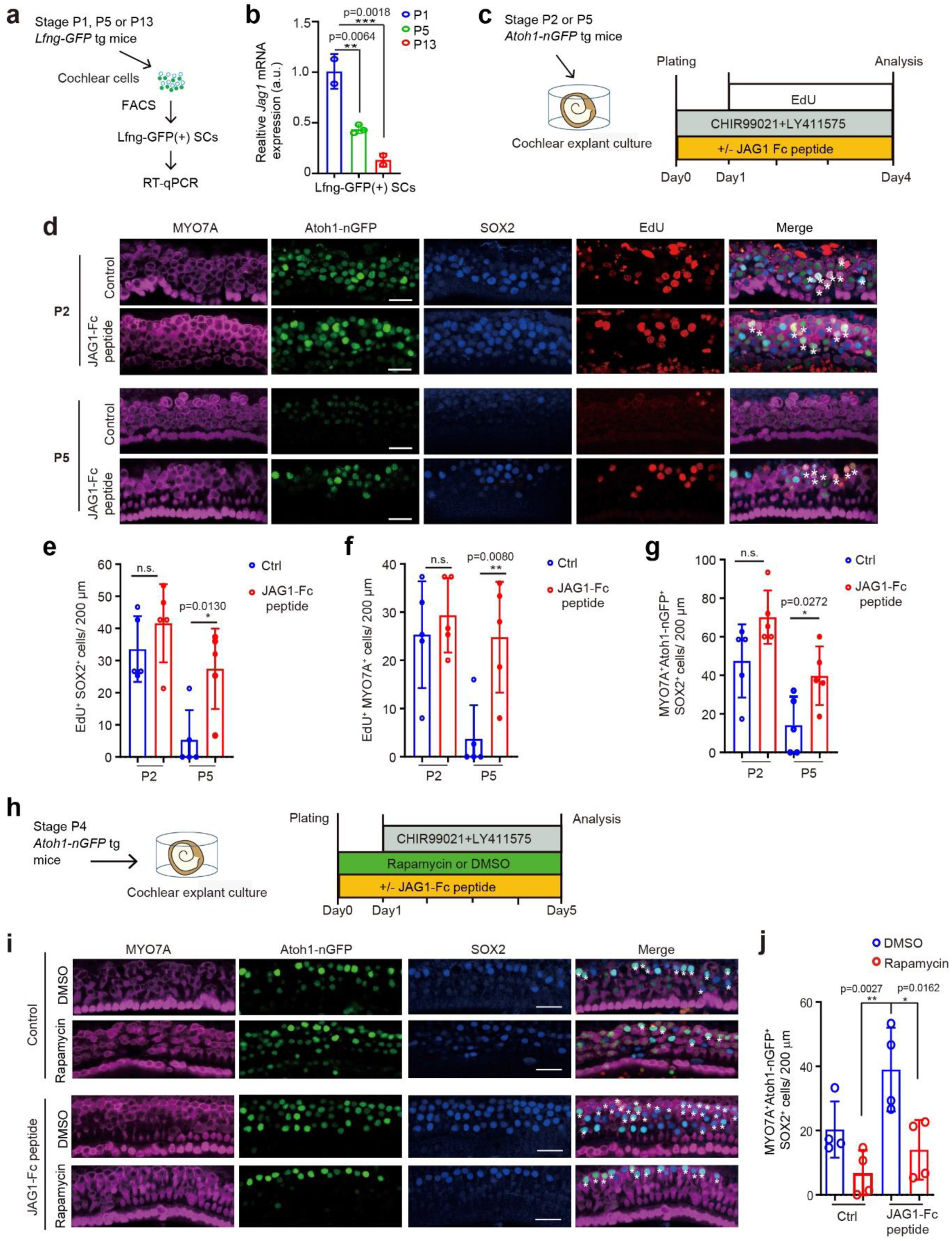
JAG1-peptide treatment boosts HC formation in cochlear explants in a stage and mTOR-dependent manner. **a** Experimental scheme for isolating Lfng-GFP^(+)^ cochlear SCs used in (**b**). **b** RT-qPCR analysis of *Jag1* mRNA expression in stage P1, P5 and P13 cochlear SCs. **c** Experimental scheme for (**d-g**). Cochlear explants from P2 or P5 *Atoh1-nGFP* tg mice were cultured with JAG1-Fc peptide or without JAG1-Fc peptide (control) in the presence of CHIR99021 and LY411575 for 4 days. EdU was added on day 1. **d** Representative confocal images of HC layer taken at the cochlear mid-apical turn following culture. Newly formed HCs (white asterisks) are labeled by MYO7A (magenta) and SOX2 (blue) immuno-staining and Atoh1-nGFP (green) expression. EdU (red) labels actively and previously dividing cells. Scale bars=25µm. **e-g** Quantification of SC proliferation (EdU^+^SOX2^+^) (**e)**, mitotic HC formation (EdU^+^MYO7A^+^) (**f**), and total number of newly formed HCs (MYO7A^+^SOX2^+^Atoh1-nGFP^+^) (**g**) for **d** (graphed are individual data points and mean ± SD, n=5 animals per group). **h** Experimental scheme for **(i-j**). Cochlear explants from stage P4 *Atoh1-nGFP* tg mice were cultured with JAG1-Fc peptide or without JAG1-Fc peptide (control) in the presence of rapamycin (4 ng/mL) or DMSO (0.05%, vehicle control) for 5 days. CHIR99021 and LY411575 were added on day 1 to induce SC-to-HC conversion. **i** Representative confocal images of the HC layer taken at the cochlear mid-apical turn following culture. Newly formed HCs are marked by MYO7A (magenta) and SOX2 (blue) immuno-staining and Atoh1-nGFP (green) expression. Scale bars=25µm. **j** Quantification of newly formed HCs in (**i**) (graphed are individual data points and mean ± SD, n = 4 animals per group, two independent experiments). Two-way ANOVA with Tukey’s correction was used to calculate *P* values. \**P* ≤ 0.05 and \*\**P*<0.01. *P* values >0.05 were considered not significant n.s.

To determine whether increasing JAG1/Notch signaling restores the mitotic and or HC-forming capacity of cochlear SCs, we isolated cochlear tissue from stage P2 and P5 Atoh1-GFP transgenic mice and cultured them as cochlear explants with JAG1-Fc peptide or without JAG1-Fc peptide (control) in the presence of CHIR99021 (Wnt activation) and LY411575 (GSI, Notch inhibitor) (Fig. 7c). EdU was added to the culture on day one to analyze for cell proliferation. After 4 days of culture, cochlear explants were stained for MYO7A, SOX2 and EdU and analyzed for SC proliferation (EdU^+^ SC^+^) and mitotic (EdU^+^ MYO7A^+^) and non-mitotic HC formation (MYO7A^+^ Atoh1-nGFP^+^ SOX2^+^) (Fig.7d). We found that in stage P2 cochlear explants the addition of JAG1-Fc peptide did not significantly increase cochlear SC proliferation (Fig.7 e, P2) or HC formation (Fig.7f and g, P2) compared to control. By contrast, in stage P5 cochlear explants, the presence of JAG1-Fc peptide significantly increased overall cell proliferation (Fig.7e), mitotic HC formation (Fig.7f) and the total number of newly formed HCs (39.7 new HCs /200 µm = 40% of total HCs) compared to control (14.1 new HCs/ 200 µm = 14% of total HCs) (Fig. 7e-g, P5). The newly formed HCs were localized in the outer HC region, indicating that SCs localized next to outer HCs, namely Deiter’s cells and pillar cells (see Fig.1a), produced the newly formed HCs.

To determine the source of the newly formed HCs, we conducted a lineage tracing experiment using cochlear tissue from stage P5 *Fgfr3icreER^T2^; R26^tdTomato/+^; Atoh1-nGFP* transgenic mice that received 4-OH tamoxifen at P3 to permanently mark Deiter’s cells and pillar cells as tdTomato+ cells (Supplementary Fig. 10a). One cochlea per animal received JAG1-Fc peptide at plating, the other functioned as control and to stimulate SC-to-HC conversion both JAG1-Fc treated, and control explants were cultured in the presence of CHIR99021 and LY411575. After 4 days of culture, we used Atoh1-nGFP reporter expression, which marks nascent HCs, to analyze the number of tdTomato+ SCs that formed new HCs at the mid-apex (Supplementary Fig. 10b). Our analysis revealed that in control cultures, only the most lateral Deiter’s cells (D3) responded to CHIR99021 and LY411575 and induced Atoh1-nGFP (9.6 new HCs/ 200 µm =10% of total HCs and 7% of total Deiter’s cells and pillar cells), while in JAG1-Fc peptide treated explants both Deiter’s cells and pillar cells upregulated Atoh1-nGFP expression in response to CHIR99021 and LY411575 (29.2 new HCs/ 200 µm =30% of total HCs and 21% of total Deiter’s cells and pillar cells) (Supplementary Fig. 10c).

We next determined whether mTOR signaling is required for JAG1’s enhancing effect on HC formation. To that end, we established cochlear explant cultures from stage P4 wild-type mice, pre-treated explants with or without JAG1-Fc peptide in the presence of rapamycin (mTORC1 inhibitor) or DMSO (0.05%, vehicle control) and the following day added CHIR99021 and LY411575 to the culture media to induce SC-to-HC conversion (Fig. 7h). After 4 days, we stained for EdU, MYO7A and SOX2 and quantified the number of newly formed HCs (MYO7A^+^ SOX2^+^ Atoh1-nGFP^+)^ ^(^Fig. 7i). We found that rapamycin reduced the number of new HCs that formed in JAG1-Fc peptide treated cultures by 3-fold (Fig. 7i and j), indicating that JAG1-Fc peptide treatment stimulates new HC formation at least in part through activating mTOR signaling.

We next analyzed how HC loss affects cochlear JAG1 expression. To ablate cochlear HCs *in vivo*, we used *Pou4f3^DTR/+^* transgenic mice, which allows us to ablate HCs using diphtheria toxin (DT). We injected stage P1 *Pou4f3^DTR/+^*transgenic mice with DT and 48 hours later immuno-stained their cochlear sensory epithelia for JAG1 protein. As a control, we analyzed JAG1 protein expression in stage P3 wild-type cochlear sensory epithelia (Fig. 8a, wild type, undamaged). We found that JAG1 expression in cochlear SCs and KCs was largely maintained following mild HC loss, but JAG1 expression was nearly absent from cochlear SCs following severe HC loss, while JAG1 expression in KCs was maintained. To determine whether stimulating JAG1/Notch signaling enhances the mitotic and or HC-regenerative capacity of cochlear SCs after severe HC loss, we established stage P5 wild-type cochlear explants, which received gentamicin for 15 hours and the following day were cultured with or without JAG1-Fc peptide in the presence of CHIR99021 and LY411575 (Fig. 8 b). After 5 days of culture, we stained control and JAG1-Fc peptide treated cochlear explants for EdU, MYO7A and SOX2 and quantified the number of newly formed HCs (MYO7A^+^ SOX2^+^) and EdU^+^ SOX2^+^ cells in the cochlear mid-apex and base (Fig. 8c). Our analysis revealed that at the cochlear mid-apex the presence of JAG1-Fc peptide allowed for some modest cell proliferation (5.6 EdU^+^SOX2^+^ cells = 3% of total SCs) (Fig. 8 c and d) and increased HC regeneration (42 new HCs/ 200µm =42% of total HCs) (Fig. 8 c, e) compared to control (23.6 new HCs/ 200µm =24%). At the cochlear base, Jag1-Fc peptide treatment (5.3 HCs/ 200µm =5% of total HCs) increased HC regeneration compared to the control (1.3 HCs/ 200µm =1% of total HCs), but the effect was low and not significant (Fig.8 c, f).

**Fig. 8.**
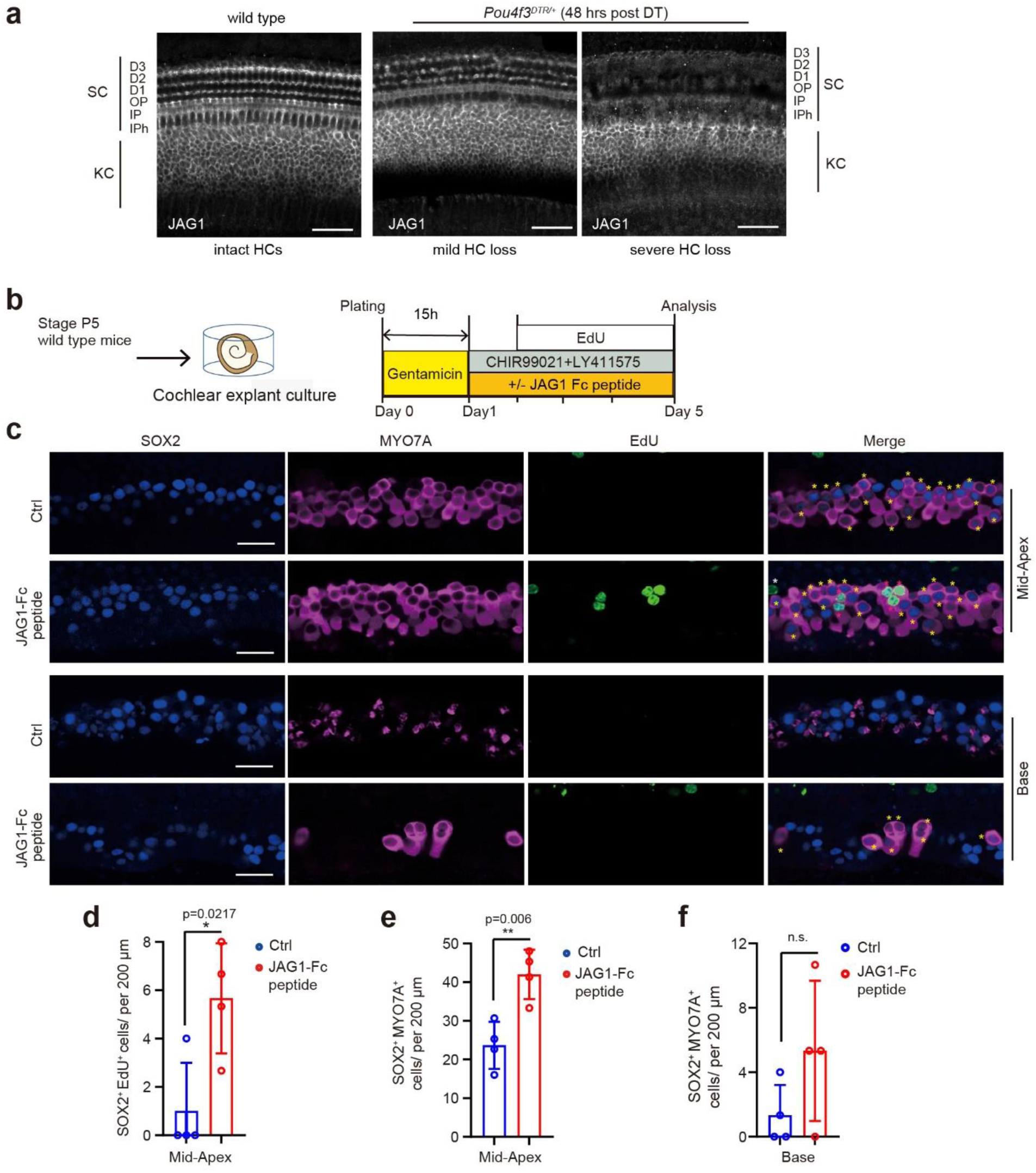
JAG1-peptide treatment enhances SC proliferation and HC regeneration in stage P5 cochlear explants. **a** Representative confocal image of JAG1 expression undamaged cochlear sensory epithelia from stage P3 wild type mice and cochlear sensory epithelia showing mild and severe HC loss from stage P3 *Pou4f3^DTR/+^* transgenic mice that received diphtheria toxin (DT) at P1. Scale bars=50µm. **(b)** Experimental scheme for (c-f). Cochlear explants were isolated from stage P5 wild type mice and after ablation of HCs with gentamicin cultured with JAG1-Fc peptide or without JAG1-Fc peptide (control) in the presence of CHIR99021 and LY411575. EdU was added to the culture media on day two to analyze cell proliferation. **c** Representative confocal images of HC layer taken at the cochlear mid-apex and base following culture. Newly formed HCs are marked by MYO7A (magenta) and SOX2 (blue) immuno-staining, indicated by yellow asterisks. EdU (green) labels actively dividing SOX2^+^ SCs (white asterisks) and previously dividing MYO7A^+^ SOX2^+^ HCs (red asterisks). Scale bars=25 µm. **d-g** Quantification of EdU^+^ SOX2^+^ cells in HC layer (HCs and SCs) (**d**), new HCs (MYO7A^+^SOX2^+^) in mid-apex (**e**) and new HCs in base (**g**) for **c** (graphed are individual data points and mean ± SD, n = 4 animals per group, two independent experiments). Two-tailed, unpaired Student’s *t* test was used to calculate *P* values. \**P* ≤ 0.05, \*\**P* <0.01. *P* values >0.05 were considered not significant n.s.

We next determined whether stimulation of JAG1/Notch signaling boosts cochlear HC formation *in vivo*. At perinatal stages, ATOH1 overexpression in cochlear SCs or KCs is highly effective in triggering their conversion into HCs. However, starting at around P7/P8, only a few new inner HCs are formed in response to ectopic ATOH1 expression^53^. To determine whether JAG1-peptide treatment can increase ATOH1-induced SC-to-HC conversion, we co-injected ATOH1-EGFP expressing AAV with JAG1-Fc peptide through the round window membrane (RWM) into the cochlea of 7-day old wild type mice. As controls, we co-injected EGFP expressing AAV with JAG1-Fc peptide and co-injected ATOH1-EGFP expressing AAV with PBS (vehicle control). We used the AAV-ie (AAV-inner ear) variant, which infects cochlear HCs, SCs, and KCs^54^. Seven days later (P14), we harvested and analyzed new HC formation in the cochlear apex and base. As expected, cochleae that received JAG1-Fc peptide and EGFP expressing control AAV (JAG1-Fc + AAV-EGFP) contained a high number of infected cells, including EGFP expressing HCs. However, none of the HCs expressed SOX2, indicating that these were pre-existing HCs. Consistent with previous reports, ATOH1 overexpression (PBS+AAV-ATOH1-EGFP) resulted in only very few new HCs (0.5 new HCs/ 200 µm= 0.5% of total HCs) within the inner HC region of the cochlear apex but close to none in the base. By contrast, cochleae that received JAG1-Fc peptide and ATOH1-EGFP expressing AAV (JAG1-Fc+ AAV-ATOH1-EGFP) contained significantly highest number of new HCs in the cochlear apex (10.5 new HCs/ 200 µm = 10.5% of total HCs) and in the cochlear base (2.5 new HCs/ 200 µm= 2.5% of total HCs) out of the three conditions (Fig.9 b, c).

**Fig. 9.**
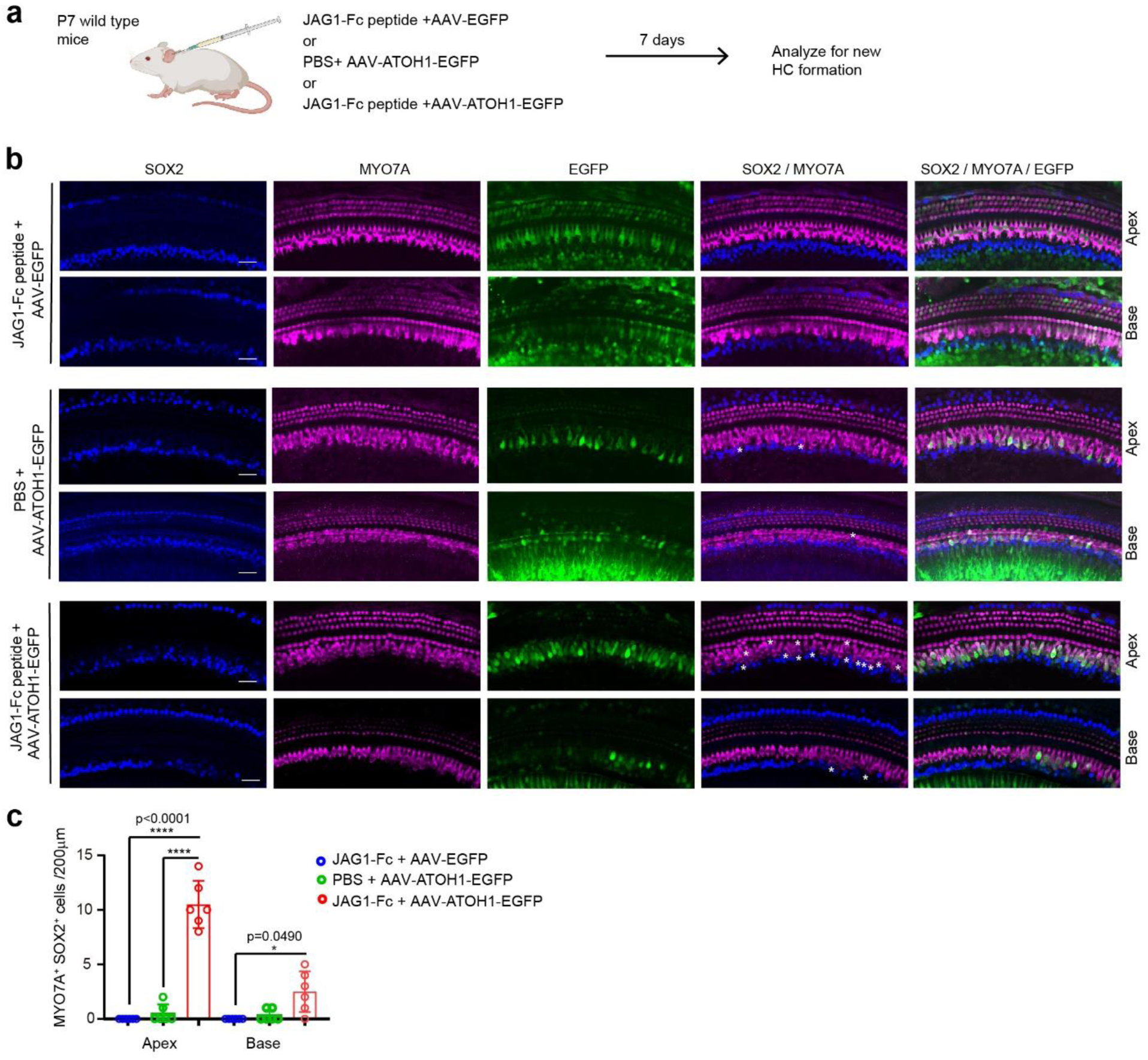
JAG1-peptide treatment enhances ATOH1-induced cochlear HC formation *in vivo*. **a** Experimental scheme. AAV expressing ATOH1 and EGFP (or EGFP only) was injected with or without JAG1-Fc peptide into the cochlea of P7 wild type mice through the round window membrane. 7 days later, injected cochleae were immuno-stained for SOX2 and MYO7A to identify newly formed HCs (MYO7A+ SOX2+). EGFP marks infected cells. **b** Representative confocal images of the HC layer at the cochlear apex and base that received JAG1-Fc peptide (JAG1-Fc peptide+ AAV-EGFP) or ATOH1 expressing AAV (PBS+ AAV-ATOH1-EGFP) (control) or both JAG1-Fc peptide and ATOH1 expressing AAV (JAG1-FC peptide+ AAV-ATOH1-EGFP). Scale bar=25 μm. **c** Quantification of infected cells that formed HCs (SOX2^+^MYO7A^+^) in the cochlear apex and base in **b** (graphed are mean ± SD, n=6 mice per group, three independent experiments). One-way ANOVA with Tukey’s correction was used to calculate P values. *P ≤ 0.05 and ****P < 0.0001.

## DISCUSSION

The safety and efficacy of GSI’s as a drug treatment for sensorineural hearing loss has been recently investigated in phase I/IIa clinical trial^55^. However, little is known about how Notch signaling operates in the HC-damaged cochlea.

The complexity of Notch ligand and receptor interactions and the limited number of cells available, paired with the dynamic and stochastic nature of HC regeneration, make it challenging to gain mechanistic insights. Notch signaling operates during cochlear HC regeneration. The recent development of organoid culture models, in which cochlear SCs can be easily manipulated, propagated, and then re-differentiated into HCs, overcomes many of these obstacles. Applying such an organoid culture model, we demonstrate two opposing functions for JAG1 in cochlear HC regeneration: (1) a negative role with JAG1 participating in HC-fate repression and (2) a positive role with JAG1 maintaining progenitor-like features of cochlear SCs, which we argue are necessary for but mitotic and non-mitotic HC regeneration.

We show that *Jag1* deficiency at perinatal stages results in a higher rate of HC formation in cochlear organoids and HC-damaged cochlear explants, indicating that JAG1 participates in HC fate repression. How does JAG1/Notch signaling restrict HC fate induction? Our transcriptomic data reveals that JAG1’s function is required to maintain the expression of *Hes1*, *HeyL,* and *Hey1* in cochlear SCs and KCs. HES1, HEYL, and HEY1 are members of the HES/HEY family of transcriptional repressors^56^, which are highly expressed in cochlear SCs ^21^ and are thought to prevent spontaneous conversion of SCs into HCs^36^. Loss of *Hes1* results in ectopic HC formation, a phenotype that is further exacerbated by the loss of *Hey1*^16,36^. However, surprisingly, *Atoh1* (*Pou4f3*) expression is only modestly elevated in *Jag1* deficient organoids, and deletion of *Jag1* in HC-depleted early postnatal cochlear explants is insufficient to trigger SC-to-HC conversion. We find that only in the presence of the GSK3 inhibitor CHIR99021 (activates Wnt signaling) does a loss of *Jag1* have an enhancing effect on cochlear HC regeneration. This is in stark contrast to the high rate of cochlear HC regeneration triggered by pharmacological inhibition of Notch signaling with GSIs ^22^ ^23^ or deletion of *Notch1*^57^, indicating a more complex role for JAG1 in cochlear HC regeneration.

Our investigation reveals that outweighing its role in HC-fate repression, JAG1’s function is essential for preserving ‘progenitor-like’ features of cochlear SCs. Previous studies have found that at perinatal stages, cochlear SCs retain some progenitor-like characteristics until perinatal stages, priming these for cell cycle re-entry and HC formation^37^. We hypothesize that maintaining or acquiring a progenitor-like state is a prerequisite for HC regeneration by SCs in general. Our hypothesis is supported by recent findings showing that in chicks and adult zebrafish, auditory and vestibular SCs de-differentiate and activate a progenitor-like state during the early phase of ‘natural’ HC regeneration^58,59^.

We find that acute deletion of *Jag1* or exposure to high dosages of GSI reduces cell cycle re-entry (organoid formation) and continued cell proliferation (organoid size), while stimulation of JAG1/Notch signaling (lentiviral *Jag1* expression, low N1ICD expression or JAG1-Fc peptide treatment) increases the frequency of cell cycle re-entry and proliferation in cochlear organoid culture. A similar pro-growth function for Notch signaling was observed when N1ICD was ectopically expressed in the murine cochlea *in vivo* at pro-sensory stages^60^.

How does JAG1/Notch signaling promote cochlear SC proliferation and HC formation? Our transcriptomic and mechanistic data indicate that JAG1/Notch1/2 signaling positively regulates PI3K-Akt-mTOR signaling in cochlear SCs and KCs. There is mounting evidence that mTOR signaling plays a critical role in cochlear SC proliferation and HC regeneration^28,61,62^. We recently showed that inhibition of mTOR activity with rapamycin reduces cochlear SC proliferation and HC formation at perinatal stages and interferes with LIN28B-induced reprogramming of cochlear SCs into progenitor-like cells^28^. Similarly, a study by Shu et al., found that reprogramming of ‘adult’ cochlear SCs with MYC and N1ICD to a developmental younger stage was sensitive to the mTOR inhibitor rapamycin and showed that MHY1485, a small molecule mTOR activator, could partially replace the need for MYC in cochlear SC reprogramming^61^. However, the above-mentioned study did not address whether Notch signaling contributes to MYC-induced mTOR activation in cochlear SCs. Our study identifies the Notch ligand JAG1 and the Notch receptors Notch1 and Notch2 as key regulators of PI3K-Akt-mTOR signaling in cochlear SCs (and KCs). We show using cochlear organoids that JAG1-Fc-peptide treatment increases PI3K-Akt-mTOR activity while loss of *Jag1,* or combined loss of *Notch1* and *Notch2* significantly reduced PI3K-Akt-mTOR activity in cochlear SCs and KCs.

Interestingly, loss of *Notch2* did not alter PI3K-Akt-mTOR activity in cochlear organoids but significantly reduced the rate of organoid formation and growth, suggesting an involvement of other JAG1-Notch2-dependent mechanisms in the regulation of cochlear organoid formation and growth. Our transcriptomic data indicates that lactate production*/*transport might be reduced in the absence of JAG1 (and potentially Notch2). A recent study found that disruption of glycolysis and lactate production reduces cochlear progenitor proliferation *in vivo*, while exogenous lactate increases cell proliferation and HC formation in cochlear organoid and explant culture^63^. Future studies are warranted to characterize a potential role for JAG1/Notch2 signaling in the regulation of cochlear glucose metabolism.

What Notch receptors mediate JAG1 function during embryonic cochlear development? We have previously shown that early embryonic cochlear deletion of *Jag1* using Emx2-Cre results in pro-sensory specification/maintenance defects with the lateral sensory domain (outer HCs and surrounding SCs) missing ^49^. By contrast, we and others have found that Notch1 deletion does not affect pro-sensory specification/maintenance^64^. Here, we tested whether previously uncharacterized Notch receptors Notch2 or Notch3 mediated JAG1’s pro-sensory function. We find that Emx2-Cre mediated *Notch2* deletion or global deletion of *Notch3* doesn’t alter cochlear HC and SC patterning, revealing that both pro-sensory specification and HC-fate repression are unaffected in these single knockout mice. Future studies using double and triple knockout approaches are required to determine the role of Notch1, Notch2, and Notch3 in pro-sensory specification and maintenance in the developing inner ear.

Studies conducted in chicken and zebrafish revealed that the strength and duration of Notch signaling are dynamically regulated during HC regeneration^3^. Initially downregulated in response to damage, Notch signaling is critical for maintaining the proper spatial and temporal pattern of SC proliferation and restricting HC-fate induction^65–68^ ^69^. Numerous studies have shown that sustained high Notch signaling blocks the ability of SCs to regenerate HCs ^70,71^.

Here, we provide evidence that too low JAG1/Notch signaling compromises the ability of murine cochlear SC to proliferate and form HCs. We show that in mice, *Jag1* mRNA expression in cochlear SCs declines more than 10-fold between stage P1 and P13 (onset of hearing) and that JAG1 protein expression in cochlear SCs greatly diminished by severe HC loss, which we hypothesize causes JAG1/Notch signaling strength to fall below the threshold needed for SCs to proliferate and form HCs. Consistent with such hypothesis are our findings that stimulation of JAG1/Notch signaling using JAG1-Fc peptide enhances SC proliferation and HC formation in cochlear explants only after the onset of cochlear maturation (stage P5) and enables ATOH1-induced HC formation by cochlear SCs in 7-day old mice *in vivo*. Recent studies have shown that ectopic ATOH1 expression in cochlear SCs is rendered ineffective due to HC-specific gene loci becoming inaccessible to ATOH1^72^. Future studies will determine whether stimulation of JAG1/Notch signaling is still effective in boosting cochlear HC formation at adult stages and whether JAG1/Notch signaling influences chromatin accessibility of HC-specific gene loci in cochlear SCs.

It is worth noting that JAG1/Notch signaling is not inhibited by GSI dosages commonly used to induce SC-to-HC conversion, suggesting that co-injection of JAG1-Fc peptide together with a low dosage of GSI may be a promising strategy to stimulate cochlear HC regeneration at later, adult stages. Taken together, our study provides mechanistic insights into the complex role of JAG1/Notch signaling in murine cochlear HC regeneration and identifies JAG1 as a novel and promising target for future therapies aimed to regenerate HCs in patients that suffered severe HC loss.

## METHODS

### Mouse breeding and genotyping

All experiments and procedures were approved by the Johns Hopkins University Institutional Animal Care and Use Committees protocol, and all experiments and procedures adhered to National Institutes of Health–approved standards. The *Atoh1-nGFP* transgenic (tg) mice were obtained from J. Johnson (University of Texas Southwestern Medical Center, Dallas). P27/GFP tg mice were obtained from Neil Segil (University of Southern California). Lfng/GFP tg mice were obtained from Nathan Heintz (Rockefeller University). *Fgfr3-iCreERT2* tg mice ^73^ were obtained from W. Richardson (University College London, UK). *Emx2^Cre/+^* mice were obtained from Shinichi Aizawa (RIKEN Kobe, Japan)^52^. *Jag1 floxed* mice were obtained from Julian Lewis (Cancer Research UK London Research Institute)^11^. *Sox2^CreER/+^* (no. 17593), *R26^rtTA*M2^* (no. 006965), *R26^LSLtdTomato/+^* (Ai14; no. 007914), *TetO-Cre* tg (no. 006234), *Notch1* floxed (f) (no. 007181), *Notch2* floxed (no. 010525) and *Notch3* −/+ (no. 010547) and *Pou4f3^DTR/+^* (no. 028673) mice were purchased from the Jackson Laboratories (Bar Harbor, ME). Mice were genotyped by PCR, as previously published. Genotyping primers are listed in Supplementary Table 4. To ablate HCs, *Pou4f3^DTR/+^* transgenic mice received a single dose of diphtheria toxin (DT) (6.25 ng/g; Sigma-Aldrich, no. D0564) by intraperitoneal injection at P1. Mice of both sexes were used in this study. All animal work was performed in accordance with the approved animal protocols from the Institutional Animal Care and Use Committees at the Johns Hopkins University School of Medicine (USA) and Xi’an Jiaotong University (China), and all efforts were made to minimize the number of mice used and their suffering.

### Viral vector and production

Lentiviral expression construct for Cre was generated using LentiCRISPRv2Cre (Addgene, no. 82415) and pCDH-EF1a-eFFly-mCherry (Addgene, no. 104833) plasmids, replacing eFFly with Cre using XbaI (NEB (New England Biolabs, no. R0145) and EagI-HF (NEB (New England Biolabs, no. R3505) restriction sites. Lentiviral expression construct for murine *Jag1* was generated by sub cloning a fragment containing 3HA-Jag1, synthesized by Sangon Biotech, into pCDH-EF1a-eFFly-mCherry (Addgene, no.104833), replacing eFFly. The pLVX-TetOne-EGFP-3HA-N1ICD plasmid, including 3HA-N1ICD fragment, was constructed using plasmid pLVX-TetO-EGFP (FENGHUISHEWU, no. BR727), linearized with EcoRI-HF (NEB (New England Biolabs, no. R3101S). All plasmids were constructed using MonClone Hi-Fusion Cloning Mix V2 (Monad, no. MC40101M). Lentiviral packaging was as conducted as previously described ^29^.

For *Notch1* and *Notch2* knockdown experiments, the plasmids for constructing the shRNAs were based on the pLV[shRNA]-mCherry/Puro-U6>Scramble shRNA#l plasmid (VectorBuilder, vector ID:VB900082.4613xkq).The sequence of shRNAs are as follows: scr-shRNA-mCherry, 5′-CCT AAG GTT AAG TCG CCC TCG-3′; Notch1-shRNA1-mCherry, 5’-ACG GCG TGA ATA CCT ACA ATT −3’; Notch1-shRNA2-mCherry,5’-CCG CTG TGA GAT TGA TGT TAA-3’; Notch1-shRNA3-mCherry, 5’-GCC AGG TTA TGA AGG TGT ATA-3’; Notch2-shRNA1-mCherry, 5’-CAC ACC AAC TTG CGC ATT AAA-3’; Notch2-shRNA2-mCherry, 5’-CGA GAC CCA GTA CAG TGA AAT −3’; Notch2-shRNA3-mChery, 5-TTC GCC TCC TGG ACG AGT ATA-3. For packaging Notch1-shRNAs and Notch2-shRNA, 293T cells are transfected with 20μg of transfer plasmid,2μg of pMDLg/pRRE (Addgene, catalog no. 12251), 2μg of pRSV-Rev (Addgene, catalog no.12253), and 2μg of pCMV-VSV-G (Addgene, catalog no.8454), respectively. After 48 hours, the supernatant was harvested and centrifuged with 25,000 RFP at 4’C for 2.5 hours, and the lentiviral particles were resuspended in DMEM/F12 to get high titer virus. The pAAV-CAG-EGFP-2A-Atoh1-3xFLAG-WPRE (7×10^12^ viral particles per mL) and pAAV-CAG-EGFP-P2A-3xFLAG-WPRE (7×10^12^ viral particles per ml) were purchased from OBiO Technology. The AAV particles were produced with AAV packaging plasmid that expressed the AAV-inner ear (AAV-ie) variant^54^.

### Fluorescence-activated cell sorting

For isolating p27-GFP^(+)^ SCs and p27-GFP^(-)^ KCs, cochlear epithelial cells from stage P2 *TetO-Cre; R26 ^rtTA*M2^*; *Jag1^f/f^; p27-GFP* transgenic animals and control littermates were used as starting material. For isolating Lfng-GFP^(+)^ SCs, micro-dissected cochlear tissue from stage P1, P2, P5 and P13 *Lfng-GFP* transgenic mice was used as starting material. Tissue was incubated in TrypLE solution (Thermo Fisher Scientific, no. 2604013), triturated, and filtered through a 35-μm filter. Resulting single cells were resuspended in expansion medium, and after incubation with propidium iodide, sorted on a MoFlo Legacy sorter with 100-μm nozzle tip.

### Organoid culture

Organoid cultures were established as previously described^28^. Briefly, dissociated cochlear epithelial cells or FACS-purified p27-GFP^(+)^ cochlear SCs or p27-GFP^(-)^ KCs from stage P2 mice were used to establish organoid cultures. In a subset of experiments cochlear epithelial cells or FACS-purified Lfng-GFP^(+)^ cochlear SCs were mixed with lentiviral particles and centrifuged at 600 g for 30 minutes prior plating. Otherwise, cells were immediately resuspended in expansion medium and mixed 1:1 with Matrigel (Corning, no. 356231), into pre-warmed four-well plates (CELLTREAT, no. 229103) and cultured in expansion media [DMEM/F12 (Corning, no. 10–092-CV), N-2 supplement (1X, ThermoFisher, no. 17502048), B-27 supplement (1X, ThermoFisher, no. 12587010), EGF (50 ng/mL, Sigma-Aldrich, no. SRP3196), FGF2 (50 ng/mL, ThermoFisher, no. PHG0264), CHIR99021 (3 μM, Sigma-Aldrich, no. SML1046), VPA (1 mM, Sigma-Aldrich, no. P4543), 616452 (2 μM, Sigma-Aldrich, no. 446859-33-2) and penicillin (100 U/mL, Sigma-Aldrich, no. P3032) to stimulate organoid formation and growth. To induce HC formation organoids were cultured in differentiation medium [DMEM/F12, N2 (1X), B27 (1X), CHIR99021 (3 μM)]. Doxycycline hyclate (Sigma-Aldrich, no. D9891) was used at 10 μg/mL if not otherwise stated. 4-hydroxy tamoxifen (Sigma-Aldrich, no. H7904) was used at 20 ng/mL. Human Jagged 1 Fc Chimera (R&D Systems, no. 1277 JG) was used at 50 ng/mL if not otherwise stated. LY411575 (Sigma-Aldrich, no. SML0506) was used as indicated. Culture media was changed every other day. Control and experimental samples were included in each independent experiment.

### Quantification of organoid forming efficiency, organoid diameter

Low-power bright-field and fluorescent images of organoid cultures were captured with an Axiovert 200 microscope using 5x and 10x objectives (Carl Zeiss Microscopy). To calculate organoid forming efficiency, the total number of organoids per culture was counted and values were normalized to the total number of cells plated. To calculate organoid size, the diameter of organoids in three to four randomly chosen fields was measured per culture using ImageJ (https://imagej.nih.gov/ij/) and the average value reported as individual data point. For each genotype and or treatment, a minimum of three biological independent organoid cultures were established and analyzed. At a minimum, two independent experiments were conducted and analyzed.

### Explant culture

Cochlear sensory epithelia including innervating neurons (cochlear explants) from stages P2-P5 mice were isolated by microdissection, and explants were cultured on SPI-Pore membrane filters (Structure Probe, no. E1013-MB) in DMEM/F12 containing 1x N-2, EGF (5 ng/mL) and penicillin (100 U/ml; Sigma-Aldrich, no. P3032). Gentamicin sulfate (100 μg/ml; Sigma-Aldrich, no. G1272) was used to ablate HCs. Recombinant human Jagged1-Fc peptide (50 ng/mL) was used to stimulate JAG1/Notch signaling. CHIR99021 (3 μM) and LY411575 (5 μM; Sigma-Aldrich, no. SML0506) were used to induce SC-to-HC conversion. To block mTORC1 activity, organoids received rapamycin (4 ng/mL; Sigma-Aldrich, no. R0395). DMSO (0.05%) served as vehicle control. Culture media was changed every other day. For each animal, two cochlear explant cultures were established, and per genotype and condition, at least three cultures obtained from different animals were analyzed. Control and experimental samples were included in each independent experiment.

### In vivo JAG1-Fc peptide experiment

Wild type stage P7 mice were anesthetized by placing them on ice for 2-3 minutes. Once anesthetized, mice were placed under the microscope and an incision was made below the left ear. The surrounding tissue and fat were carefully removed with forceps to expose the round window membrane of the cochlea. 500 nL JAG1-Fc peptide (50μg/mL) dissolved in PBS or 500 nL PBS was injected together with 500 nL AAV-EGFP or 500 nL AAV-ATOH1-EGFP at the speed of 20nL/sec by a pressure controlled motorized micro injector (RWD, R-480). An equal volume of AAV was used and the total volume for each injection was 1μl. After the injection, the skin incision was closed using veterinary tissue adhesive (Millpledge Ltd, UK). Pups were subsequently returned to their mother for continued nursing.

### RNA extraction, RT-qPCR and TaqMan assay

Lfng-GFP^(+)^ cochlear SCs at from stage P1, P5 and P13 Lfng-GFP transgenic mice were isolated by FACS as previously described^29^. Organoids were harvested using Cell Recovery Solution. Total RNA from acutely isolated SCs or cultured cochlear organoids/explants was extracted using the miRNeasy Micro Kit. MRNA was reverse transcribed into cDNA using the iScript cDNA synthesis kit. Q-PCR was performed on a CFX-Connect Real Time PCR Detection System using SYBR Green Master Mix reagent. Gene-specific primers used are listed in Supplementary Table 5.

### RNA sequencing and data analysis

For each condition, three independent cultures from three animals were established. All the samples were processed using Illumina’s TruSeq stranded Total RNA kit, per manufacturer’s recommendations, using the UDI indexes. The samples were sequenced on the NovaSeq 6000, paired end, 2×50 base pair reads. Kallisto (v0.46.1) was used to pseudo-align reads to the reference mouse transcriptome and to quantify transcript abundance. The transcriptome index was built using the Ensembl Mus musculus v96 transcriptome. The companion analysis tool sleuth was used to identify differentially expressed genes (DEGs). We performed a Wald test to produce a list of significant DEGs between *Jag1* KO expressing sample and the control. These lists were then represented graphically using sleuth along with pheatmap and ggplot2 packages in R v1.3.1093. Gene identifier conversion, gene annotation, and enrichment analysis was conducted using Metascape^42^.

### Immunohistochemistry

Cochlear organoids/explants were fixed in 4% paraformaldehyde for 30 minutes, permeabilized and blocked with 0.25% Triton X-100/ 10% fetal bovine serum for 30 minutes, and immuno-stained as previously described^28^. Antibodies are listed in Supplementary table S6.

### Cell proliferation

EdU (ThermoFisher Scientific, no. C10338) was added to culture medium at a final concentration of 3 µM as indicated after which cochlear organoids /explants were harvested and processed for EdU detection using the Click-iTPlus EdU Cell Proliferation Kit (ThermoFisher Scientific, no. C10637 and C10638) following the manufacturer’s recommendations.

### Quantification of cell proliferation and HC formation

High-power confocal single-plane and z-stack images of fluorescently immunolabeled organoids and explants were taken with 40× objective using LSM 700 confocal microscope (Zeiss Microscopy). Explants: For each explant three high power confocal images of the HC-layer were taken and used to determine the average number of EdU^+^ cells and MYO7A^+^SOX2^+^ HCs per length unit. Orientation of shown cochlear sensory epithelium is such that medial inner HC region is closest to the bottom edge and lateral outer HC region is closest to the top edge of shown images. HC formation and cell proliferation was quantified in the cochlear mid-apex if not otherwise stated. At least three biological independent cultures per group (treatment/genotype) were analyzed and reported as individual data points. Organoids: For each culture, low power and high-power confocal images of processed organoids were taken and used to calculate the average percentage of EdU^+^ cells and MYO7A^+^SOX2^+^ HCs per organoid. At least three biological independent cultures per group (treatment/genotype) were analyzed and reported as individual data points.

### Protein lysis and immunoblotting

Cochlear organoids were lysed with RIPA buffer (Sigma-Aldrich, no. R0278) supplemented with protease inhibitor (Sigma-Aldrich, no.11697498001), phosphatase Inhibitor cocktail 2 (Sigma-Aldrich, no. P5726) and phosphatase inhibitor cocktail 3 (Sigma-Aldrich, no. P0044). Protein separation and immunoblots were conducted as previously described^28^. The resulting chemiluminescence was captured using X-ray films or digitally using the LI-COR Odyssey imaging system. ImageJ (https://imagej.nih.gov/ij/) was used to quantify the protein levels by measuring the relative density of bands. The antibodies used are listed in Supplementary Table S7

### Statistical analysis

All results were confirmed by at least two independent experiments. Control and experimental samples were included in each independent experiment. The sample size (n) represents the number of animals analyzed per group if not otherwise stated. Animals (biological replicates) were allocated into control or experimental groups based on genotype and/or type of treatment. Derived statistics correspond to analysis of averaged values across biological replicates, and not pooled technically and biological replicates. To avoid bias, masking was used during data analysis. Data was analyzed using GraphPad Prism 8.0. Relevant information for each experiment, including sample size, statistical tests, and reported *P* values, are found in the legend corresponding to each figure. In all cases, *P* values ≤0.05 were considered significant, and error bars represent standard deviation (SD).

## ACKNOWLEDGMENT

We thank the members of the Doetzlhofer Laboratory for their help and advice throughout this study.

## Author’s contributions

Conceptualization: A.D., X.-J.L.

Methodology: A.D., X.-J.L, C.M., Investigation: X.-J.L., C.M., L.L., W.-Y.Z., E.C.

Supervision: A.D., X.-J.L Writing—original draft: A.D., X.-J.L.

Writing—review & editing: A.D., X.-J.L., C.M

## Data availability

RNA sequencing data have been deposited in the Gene Expression Omnibus data repository under accession number: GSE266157. All data supporting the findings of this study are available within the main manuscript and its supplementary information. Source Data are provided for this study.

## AUTHORS CONTRIBUTIONS

X.-J.L. and A.D. designed the research; X.-J.L., C.M., L.L., W.-Y.Z, and E.C. performed the research; X.-J.L., C.M., and A.D. analyzed the data; and X.-J.L. and A.D. wrote and critically reviewed the manuscript.

