## Supplementary Figures for "The Notch ligand Jagged1 plays a dual role in cochlear hair cell regeneration"

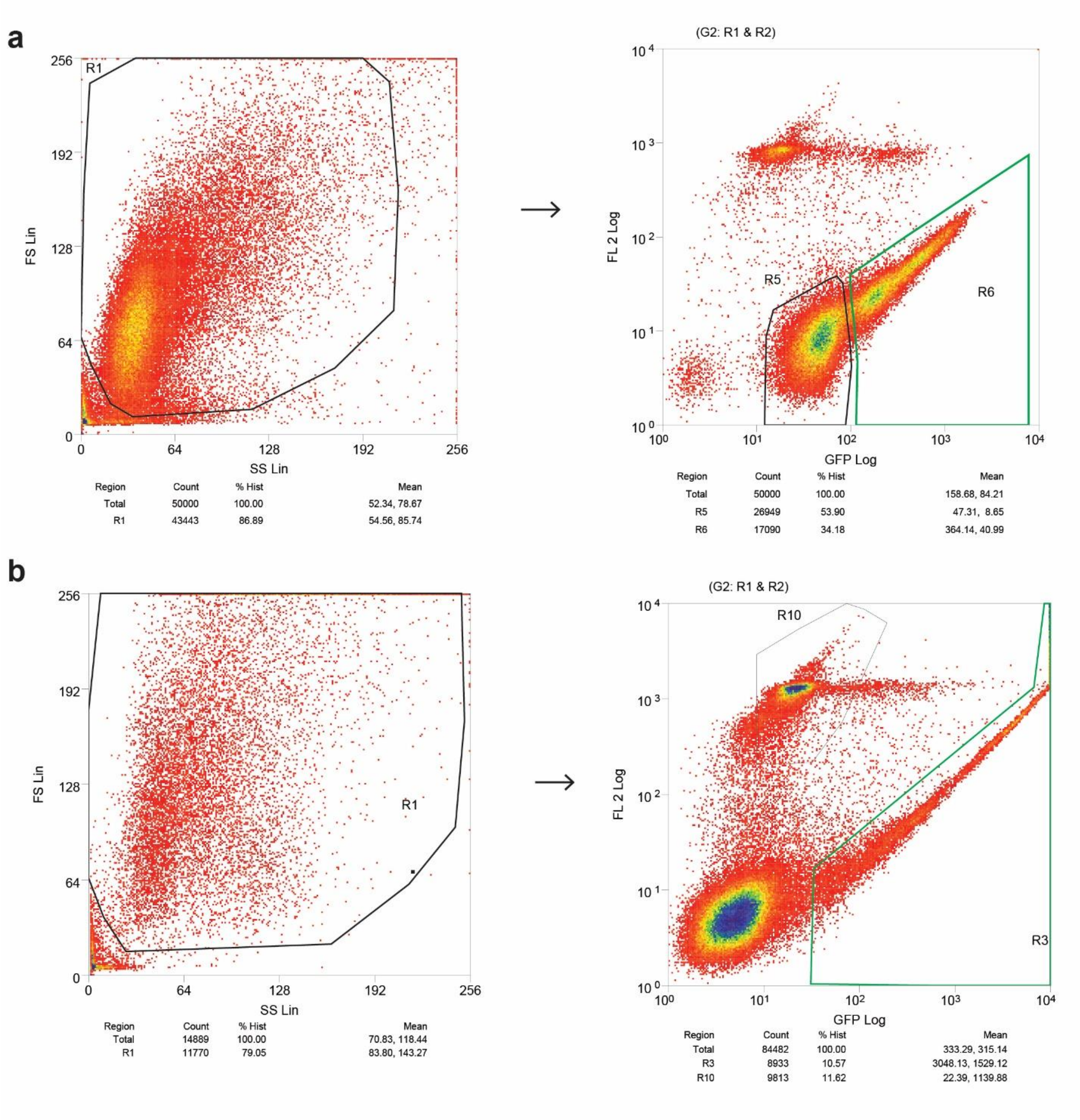


**Supplementary Fig.1. Fluorescent Activated Cell Sorting (FACS) of cochlear SCs using p27-GFP and Lfng-GFP.** Shown are representative forward scatter (FS) /side scatter (SS) plots (left) and FACS plots (right) of GFP (x-axis) versus FL2 (y-axis, PI). Before sorting, Propidium Iodide (PI) was added to cell suspension to visualize dead cells. **a** Gating strategy for collection of p27-GFP(+) SCs and p27-GFP(-) KCs. Cochlear epithelial cells from stage P2 p27-GFP transgenic mice were gated on a live cell population using an FS/SS gate (R1). R6 (green box) indicates gating for GFP(+) SCs, and R5 (black box) indicates gating for GFP(-) KCs. **b** Gating strategy for collection of Lfng-GFP(+) SCs. Stage P1 Lfng-GFP transgenic cochlear cells were gated on a live cell population using an FS/SS gate (R1). R3 (green box) indicates gating for GFP(+) SCs.

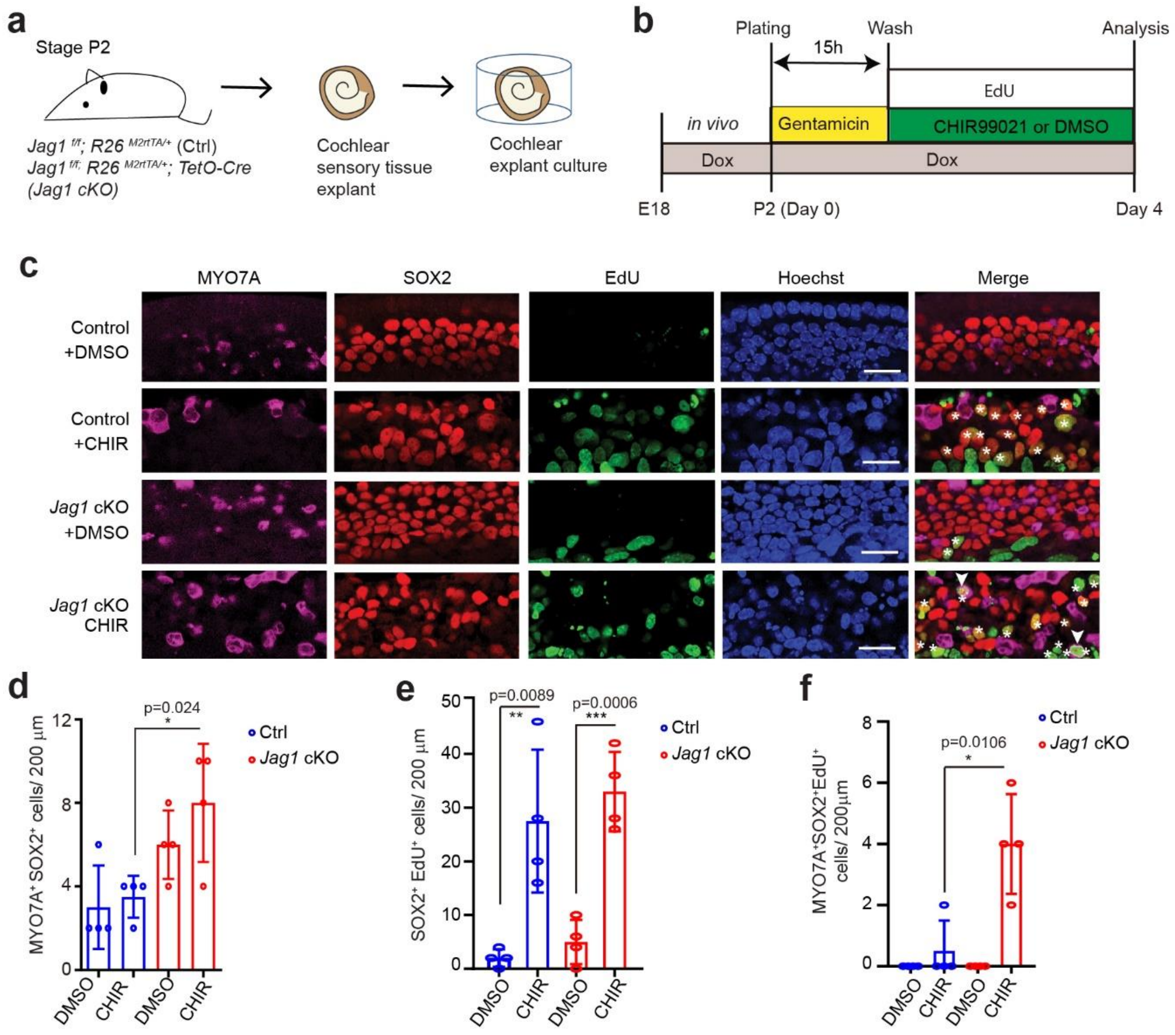


**Supplementary Fig. 2. Loss of *Jag1* in the HC-damaged cochlear explants enhances HC regeneration induced by Wnt activation. a** Experimental scheme. Cochlear explant cultures were established from stage P2 *Jag1* conditional knockout (cKO) (T*etO-cre tg/+; R26*$^{M2rtTA/+}$*; Jag1*$^{f/f}$) mice and control littermates (*R26*$^{M2rtTA/+}$*; Jag1*$^{f/f}$). **b** Experimental timeline. Pregnant dams were fed doxycycline (dox) containing food starting at E18.5, and dox was present throughout the duration of the culture. Gentamicin was added to the culture media at plating and removed the next day. EdU was present throughout the duration of the culture. CHIR99021 (3 µM) or DMSO (vehicle control) was added the next day (day 1). The culture media was replenished every day. **c** Shown are representative confocal images of MYO7A (magenta), SOX2 (red), EdU (green), Hoechst (blue) in the mid-apical turn of control and *Jag1* CKO mice cultured with DMSO or CHIR99021 (CHIR). White asterisks mark proliferating SCs (EdU+SOX2+), and white triangles mark newly formed HCs (MYO7A+; SOX2+). Scale bars=25µm. **d-f** Quantification of HC formation (**d**), SC proliferation (**e**), and mitotic HC formation (**f**) in (**c**) (graphed are average values for each animal and their mean± SD, n=4 animals per group, two-way ANOVA with Tukey's correction was used to calculate p-values). *$P \leq 0.05$, **$P < 0.01$, ***$P < 0.001$.

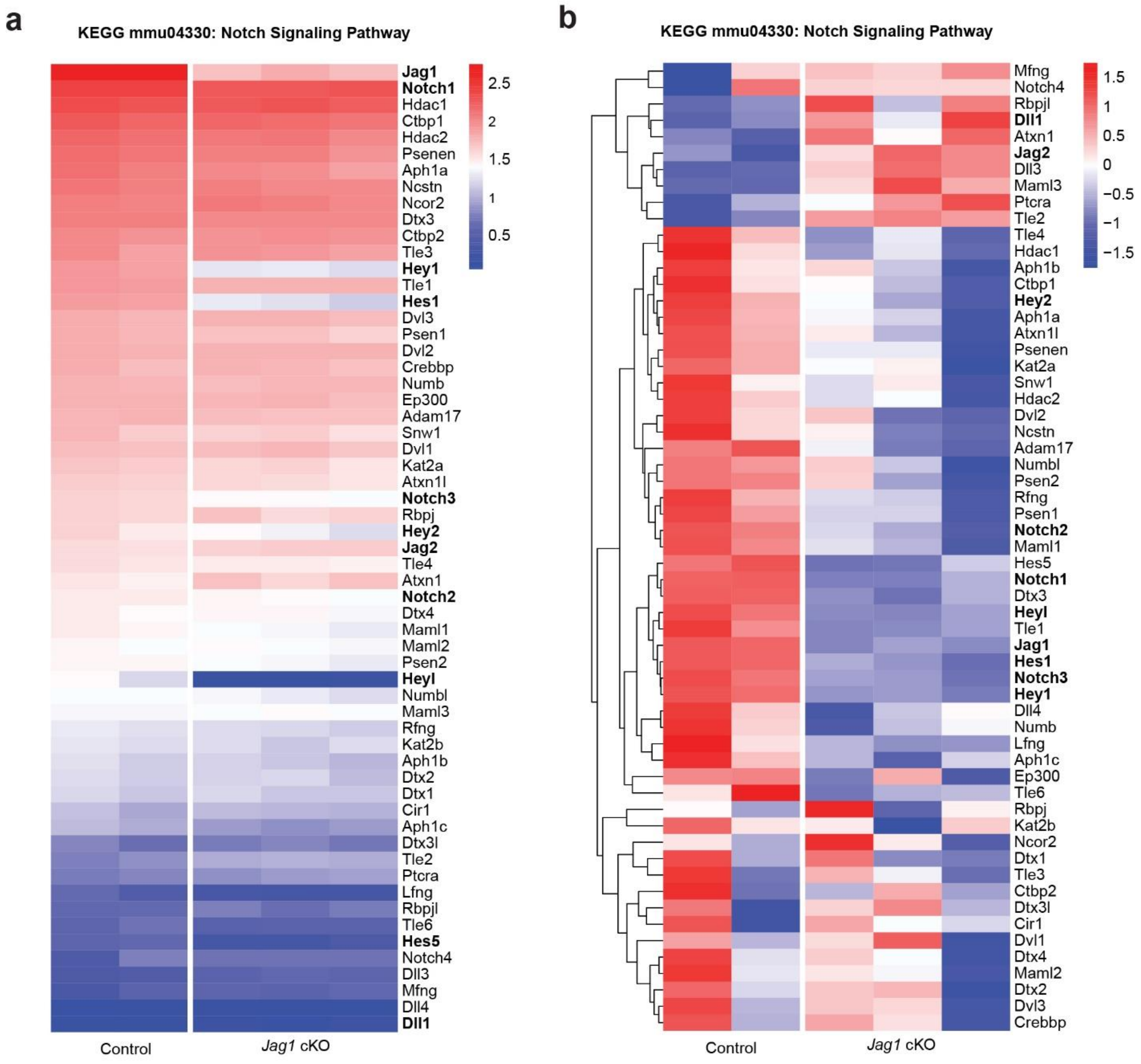


**Supplementary Fig. 3. Regulation and expression of genes associated with Notch signaling pathway in control and *Jag1* cKO organoids.** Heatmaps display the abundance (a) and relative expression (b) of gene transcripts associated with KEGG mmu04330: Notch Signaling Pathway in control and *Jag1* cKO condition. Bold font was used for ligands, receptors, and effectors of canonical Notch signaling. a Counts have been color-coded to indicate high (red) and low abundant (blue) transcripts in *Jag1* cKO and control conditions. b Row-normalized counts have been color-coded to show relative increases (red) or decreases (blue) in transcript abundance in the *Jag1* cKO condition compared to control, and hierarchical clustering was used to group genes with similar expression patterns.

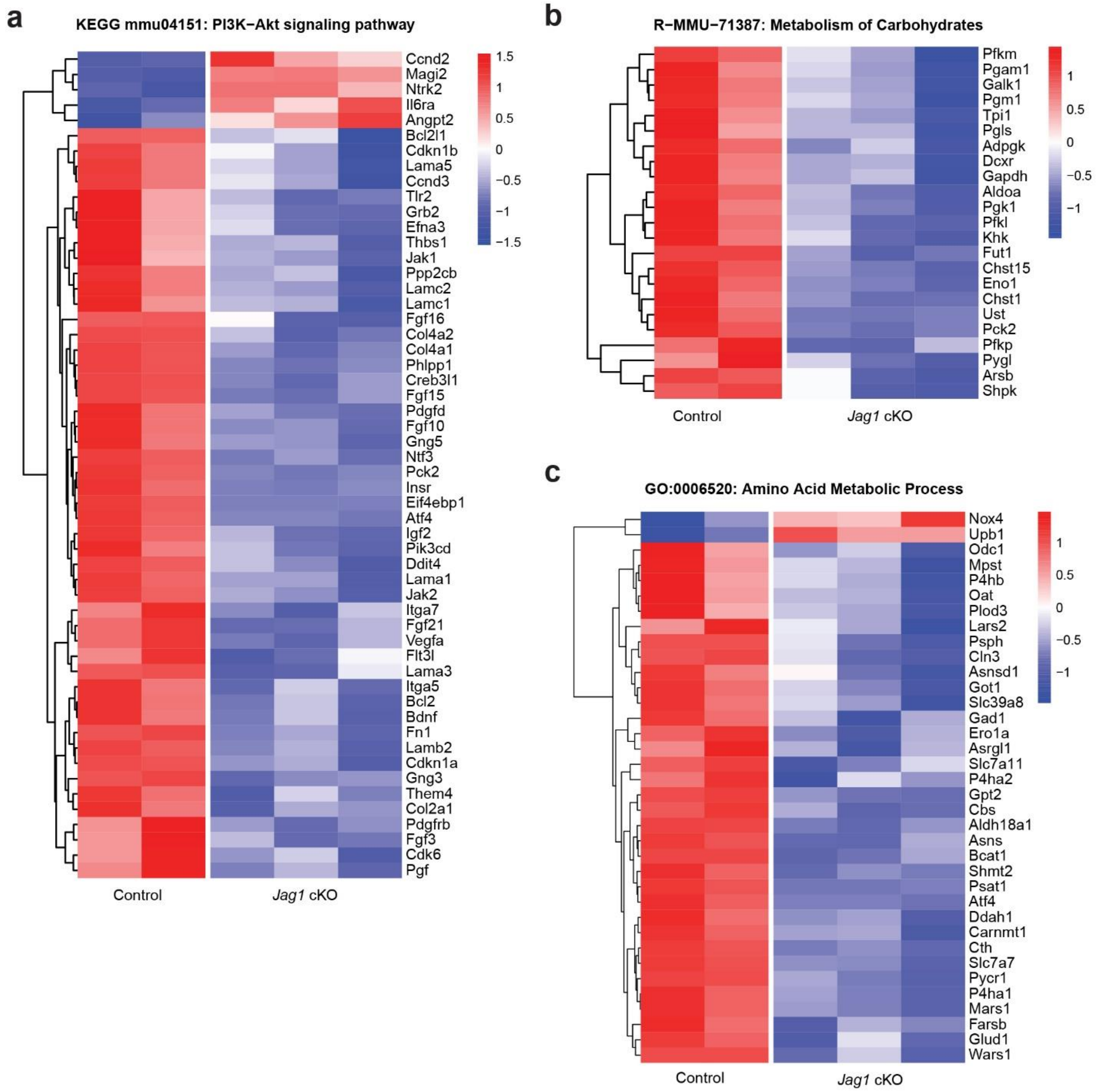


**Supplementary Fig. 4. Regulation of genes associated with PI3K-Akt signaling and metabolic pathways in control and *Jag1* cKO organoids.** Row-normalized counts were color-coded to show relative increases (red) or decreases (blue) in expression in the *Jag1* cKO condition compared to control, and hierarchical clustering was used to group genes with similar expression patterns. a Heatmap displaying differentially expressed genes (DEG) associated with KEGG mmu04151: PI3K-Akt signaling pathway. b Heatmap displaying DEG associated with R-MMU-71387 Metabolism of Carbohydrates. c Heatmap displaying DEG associated with GO: 0006520: Amino Acid Metabolic Process.

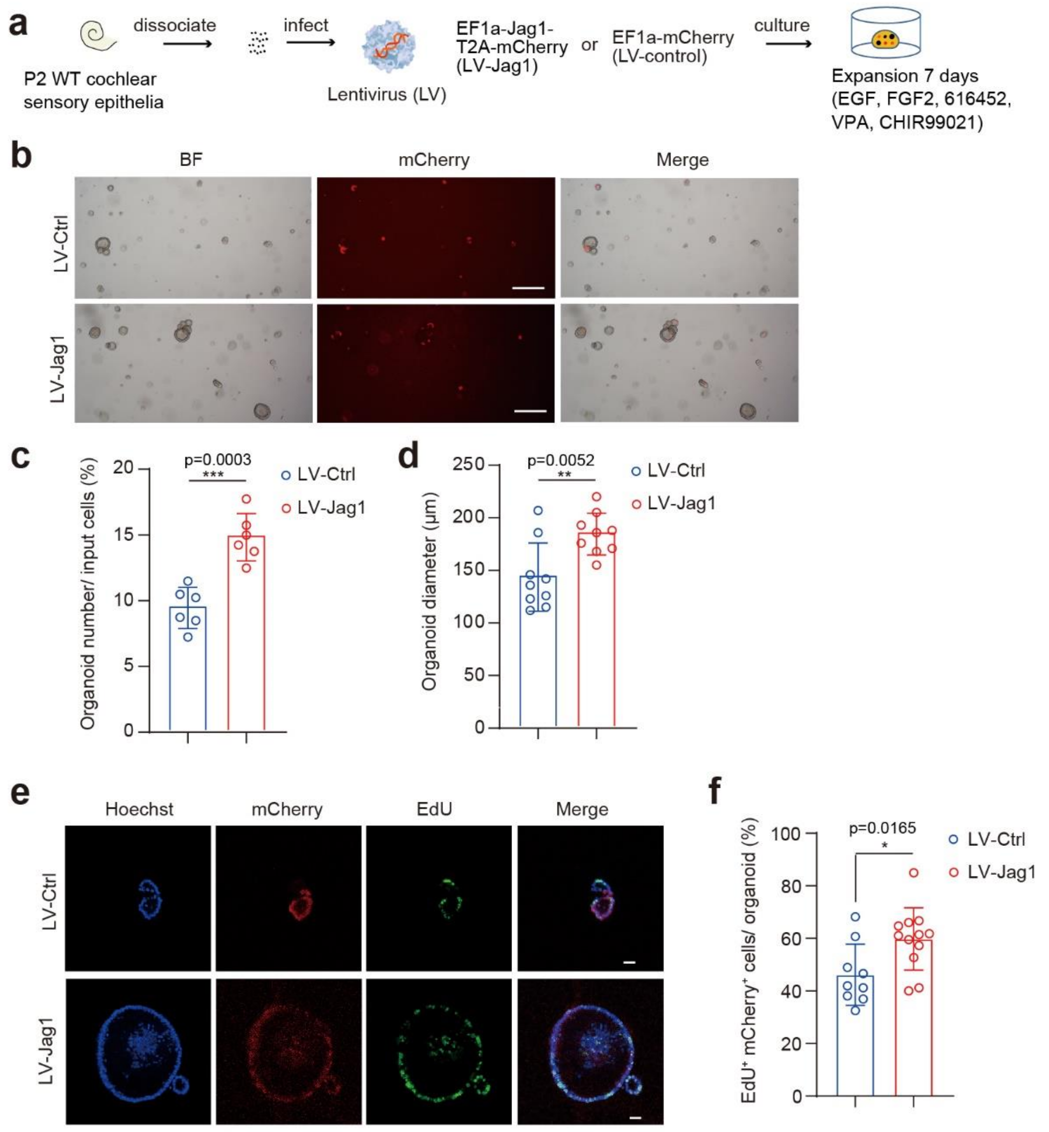


**Supplementary Fig. 5. Lentiviral overexpression of JAG1 stimulates cochlear organoid formation and growth**. **a** Experimental scheme. Cochlear epithelial cells from P2 wild-type mice were infected with mCherry expressing control or mCherry and JAG1 expressing lentivirus (LV) and expanded as organoids for 7 days. **b** Bright-field (BF) and red fluorescent (mCherry) images of organoids infected with control and JAG1 expressing lentivirus at 7 days of expansion. Scale bars=400µm. **c** organoid forming efficiency in (**b**) (graphed are individual data points and mean ± SD, n=6 independent biological replicates, two independent experiments). **d** Organoid diameter in (**b**) (graphed are individual data points and mean ± SD, n=9 independent biological replicates, three independent experiments). **e** Cell proliferation in organoids infected with control and JAG1 expressing lentivirus. A single EdU pulse was given at 7 days of expansion, and EdU incorporation (green) was analyzed 1 hour later. Hoechst labels cell nuclei (blue), and mCherry labels infected cells (red). Scale bars=25µm. **f** Percentage of EdU+ cells in (d) (graphed are individual data points and mean ± SD, n=9 independent biological replicates for organoid cultures infected with control and n=12 independent biological replicates for organoid cultures infected with JAG1 expressing lentivirus). Two-tailed, unpaired Student's *t* test was used to calculate *P* values. $^{*}P \leq 0.05$, $^{**}P<0.01$ *and* $^{***}P<0.001$.

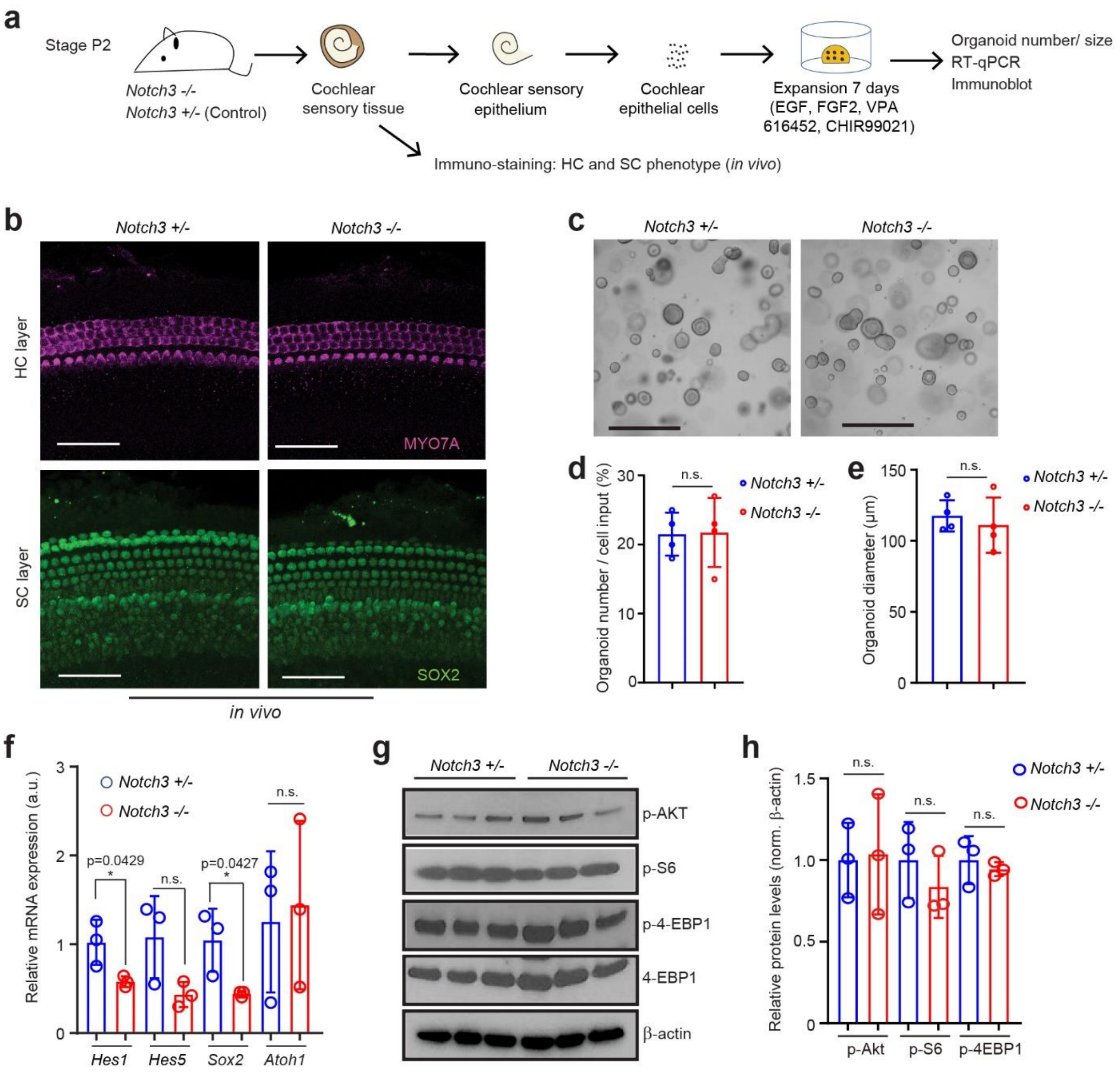


**Supplementary Fig. 6. Loss of Notch3 does not alter cochlear organoid formation or growth and does not reduce PI3K-Akt-mTOR signaling.** **a** Experimental scheme. **b** Confocal images of HC and SC layers of cochlear sensory epithelia from stage P2 *Notch3*$^{-/-}$ (knockout, KO) and *Notch3*$^{-/+}$ (control) mice. MYO7A (magenta) labels HCs, and SOX2 (green) labels SCs. Scale bars=50µm. **c** Bright field (BF) images of cochlear organoid cultures established from P2 *Notch3*$^{-/+}$ (control) and *Notch3*$^{-/-}$ (KO) mice at 7 days of expansion. Scale bars=400µm. **d-e** Organoid forming efficiency (d) and organoid diameter (e) for c (graphed are individual data points and mean ± SD, n=4 animals per group). **f** RT-qPCR of Notch target genes (*Hes1*, *Hes5,* and *Sox2*) and *Atoh1* in *Notch3*$^{-/+}$ (control) and *Notch3*$^{-/-}$ (KO) organoids at 7 days of expansion (graphed mean ± SD, n=3 animals per group). **g** Immunoblot of p-Akt, p-S6, p-4-EBP1, 4-EBP1 and β-actin proteins in cochlear organoids established with control and *Notch3* KO mice after 7 days of expansion. **h** p-S6 and p-4EBP1 protein levels normalized to β-actin for g (graphed are individual data points and mean ± SD, n=3 animals per group). Two-tailed, unpaired Student's *t* test was used to calculate P values. *P* >0.05 was deemed not significant (n.s.).

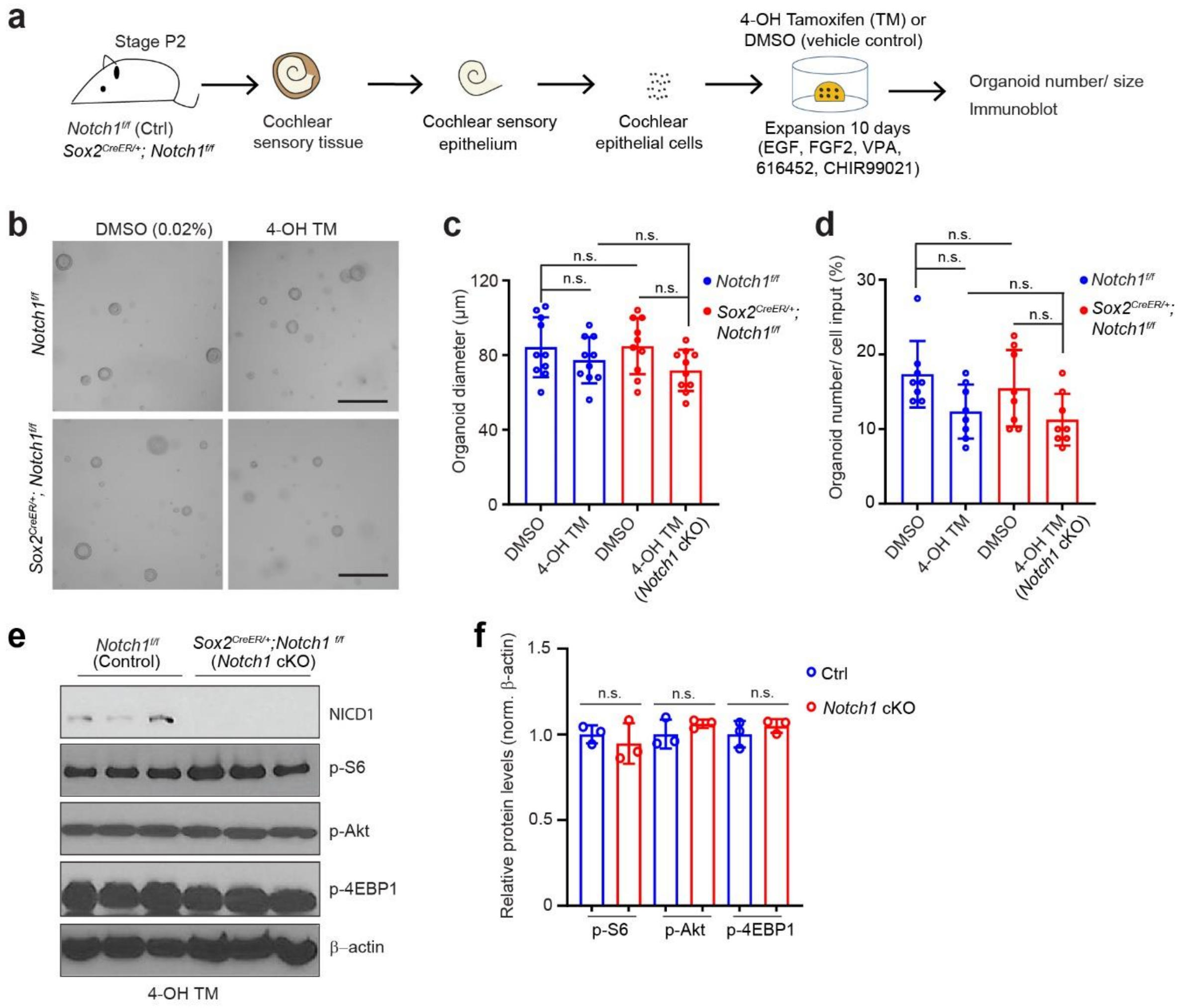


**Supplementary Fig. 7. Loss of *Notch1* does not alter cochlear organoid formation and growth, nor does loss of Notch1 reduce PI3K-Akt-mTOR signaling. a.** Experimental scheme. Organoids were established with cochlear epithelial cells from stage P2 *Notch1*$^{f/f}$ and *Sox2*$^{CreER/+}$*; Notch1*$^{f/f}$ mice. Organoids were cultured in the presence of 4-OH tamoxifen (TM) or DMSO (vehicle control). **b** Bright field (BF) images of control [(DMSO: *Notch1*$^{f/f}$), (DMSO: *Sox2*$^{CreER/+}$ *Notch1*$^{f/f}$), (4-OH TM: *Notch1*$^{f/f}$)] and *Notch1* KO organoid cultures (4-OH TM: *Sox2*$^{CreER/+}$*; Notch1*$^{f/f}$) at 10 days of expansion. Scale bars=400μm. **c** Quantification of organoid diameter in (b) (graphed are individual data points and mean ± SD, n=10 animals per group). **d** Quantification of organoid forming efficiency in (b) (graphed are individual data points and mean ± SD, n=8 animals per group). **e** Immunoblots of NICD1, p-S6, p-Akt, p-4EBP1 and β-actin proteins in *Notch1*$^{f/f}$ (control) and *Sox2*$^{CreER/+}$*; Notch1*$^{f/f}$ (*Notch1* KO) organoids at 10 days of expansion. **f** Normalized protein levels of NICD1, p-Akt, p-S6, p-4EBP1 in (e) (graphed are individual data points and mean ± SD, n=3 animals per group). Two-tailed, unpaired Student's *t* test was used to calculate *P* values. *P* >0.05 was deemed not significant (n.s.).

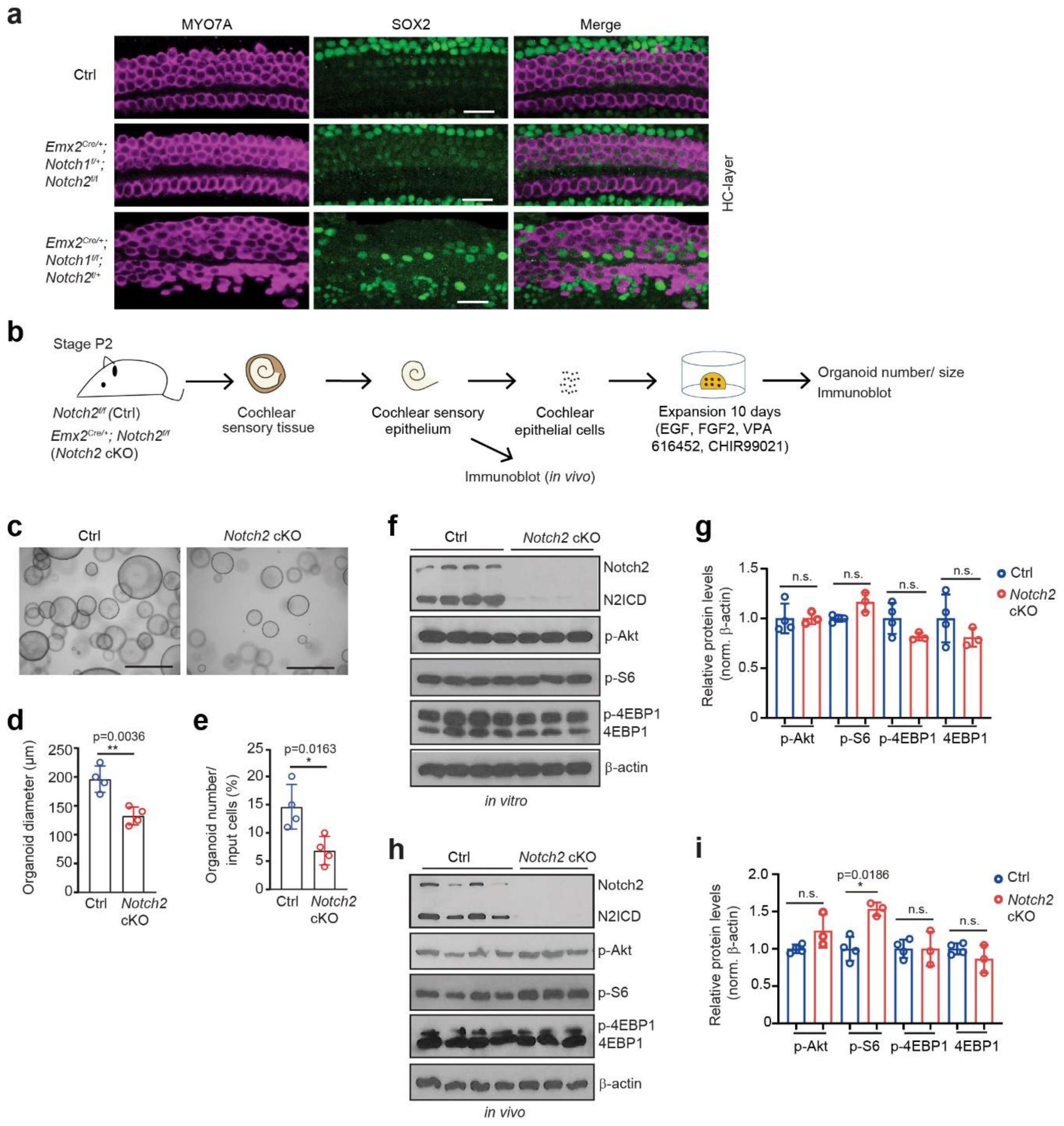


**Supplementary Fig. 8. Loss of Notch2 reduces cochlear organoid formation and growth but does not alter PI3K-Akt-mTOR signaling. a** Confocal image of cochlear sensory epithelia of *Notch2* and *Notch1* homozygous and heterozygous mutant mice (*Emx2*$^{Cre/+}$; *Notch2*$^{f/f}$; *Notch1*$^{f/+}$ and *Emx2*$^{Cre/+}$; *Notch2*$^{f/+}$; *Notch1*$^{f/f}$) and *Emx2*$^{Cre/+}$ negative control (Ctrl) littermates. MYO7A (magenta) labels HCs, and SOX2 (green) labels SCs. MYO7A$^{+}$SOX2$^{+}$ cells represent recently formed HCs. Scale bars=25µm. **b** Experimental scheme for (c-i). **c** Bright field (BF) images of P2 *Notch2*$^{fl/fl}$ (control) and *Emx2*$^{Cre/+}$*; Notch2*$^{f/f}$ *(*cKO) organoids at 10 days of expansion. Scale bars=400µm. **d-e** Organoid diameters (**d**) and organoid formation efficiency (**e**) for **c** (graphed are individual data points and mean ± SD, n=4 animals per group). **f** Immunoblot of Notch2, NICD2, p-Akt, p-S6, p-4EBP1 and β-actin proteins in cochlear organoids established with control and *Notch2* CKO mice. **g** Normalized protein levels of Notch2, NICD2, p-Akt, p-S6, p-4EBP1 in (**f**) (graphed are individual data points and mean ± SD, n=4 animals per group ). **h** Immunoblot of Notch2, NICD2, p-Akt, p-S6, p-4EBP1 and β-actin proteins in cochlear epithelial cells isolated from $Notch2^{fl/fl}$ (control) and $Emx2^{Cre/+}$ $Notch2^{fl/fl}$ *(Notch2* KO) mice at P5. **i** Normalized protein levels of Notch2, NICD2, p-Akt, p-S6, p-4EBP1 in (h) (graphed are individual data points and mean ± SD, n=4 animals for control and n=3 mice for *Notch2* KO). Two-tailed, unpaired Student's *t* test was used to calculate P values. *$P \leq 0.05$, **$P < 0.01$.

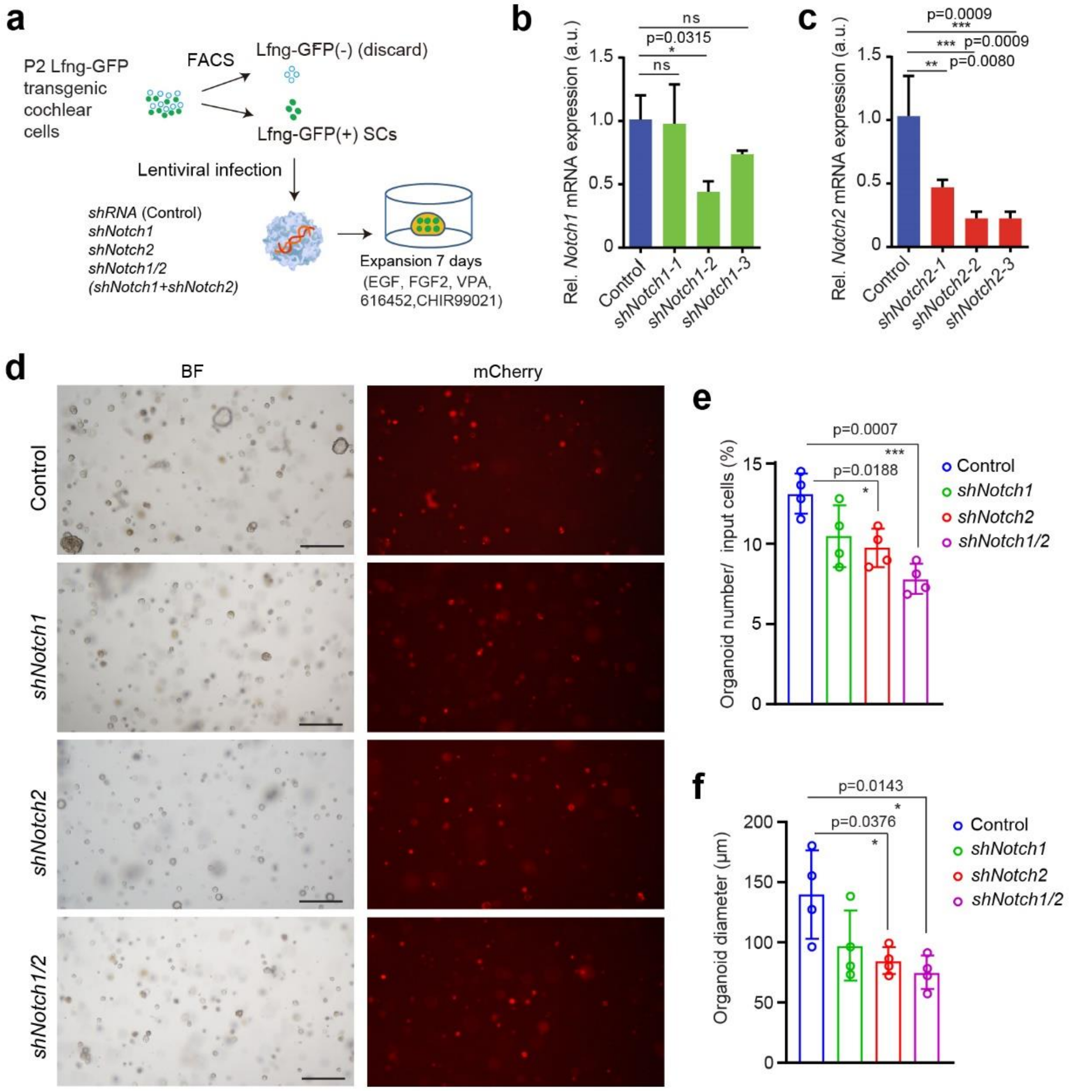


**Supplementary Fig. 9. Knockdown of *Notch1* and Notch2 in cochlear SCs inhibits organoid formation and growth.** **a** Experiment scheme. Organoid cultures were established with FACS-purified Lfng-GFP$^{(+)}$ SCs from stage P2 *Lfng-GFP* tg mice, and lentiviral particles expressing control *shRNA*, *shRNA-Notch1*, *shRNA-Notch2*, *shRNA-Notch1/2* were used to infect the Lfng-GFP$^{(+)}$ SCs before plating. **b** RT-qPCR analysis of knockdown efficiency of *shNotch1-1*, *shNotch1-2*, *shNotch1-3* (bar plots represent mean values ± SD), *shNotch1-2* was used to subsequent experiments. **c** RT-qPCR-based analysis of the knockdown efficiency of *shNotch2-1*, *shNotch2-2*, *shNotch2-3* (bar plots represent mean values ± SD), and *shNotch2-3* on its own or in combination with *shNotch1-2* was used in subsequent experiments. **d** Bright field (BF) and red fluorescence (mCherry) images of SC-derived organoid culture after knockdown *Notch1, Notch2* or *Notch1/2.* MCherry expression marks infected cells. Scale bar=400 μm. **e-f** Organoid formation efficiency (**e**) and organoid diameter (**f**) in d (graphed are individual data points and mean ± SD, n=4 independent biological replicates, from one representative experiment and two independent experiments). One-way ANOVA with Tukey's correction was used to calculate *P* values. *P ≤0.05, **P <0.01 and ***P < 0.001.

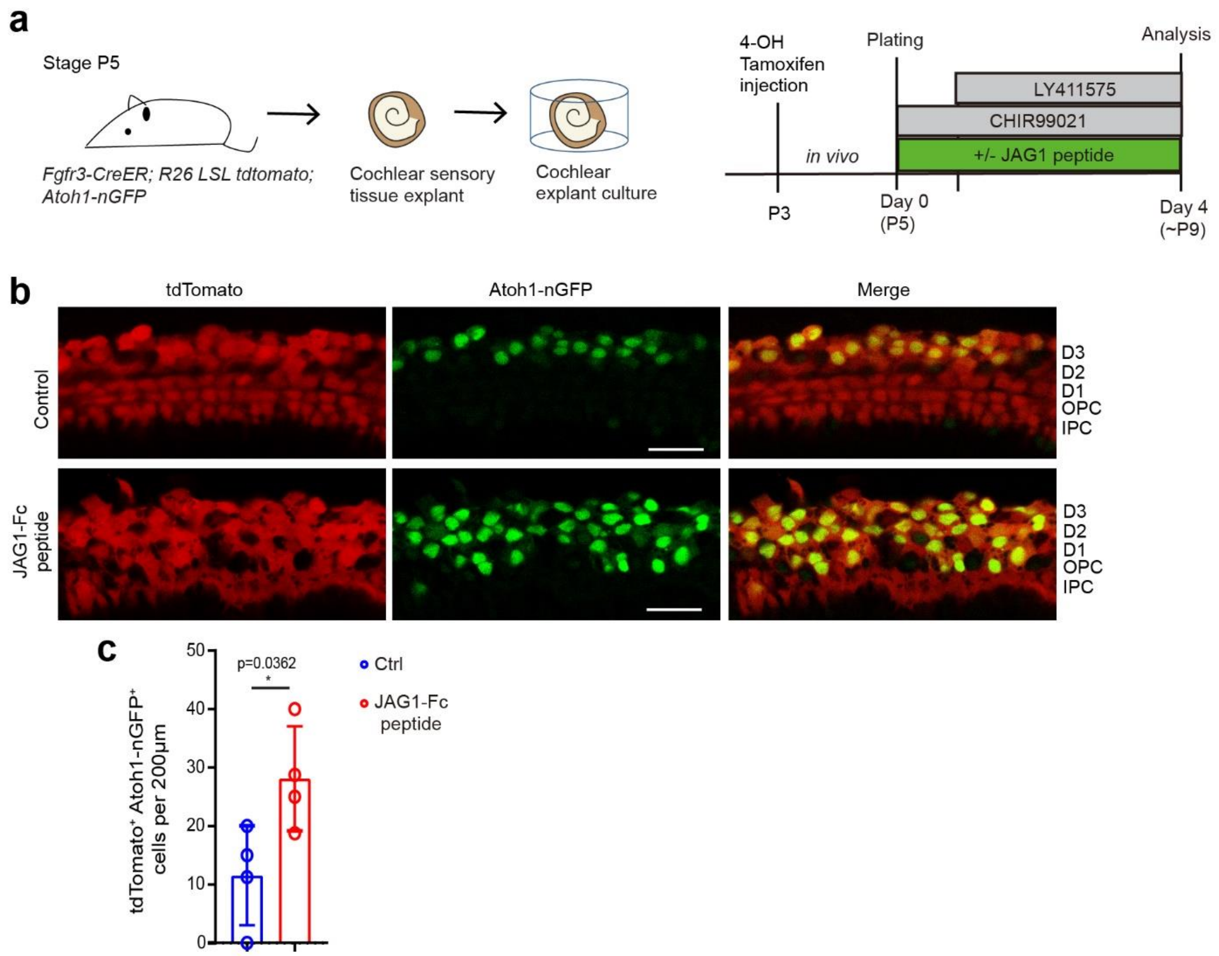


**Supplementary Fig. 10. JAG1-peptide treatment increases the rate of cochlear SC-to-HC conversion in response to Wnt activation and Notch inhibition. a** Experimental scheme. Cochlear explant cultures were derived from P5 *Atoh1-nGFP* tg; *Fgfr3icreER; R26* $^{\text{LSL-tdTomato}}$ mice, which received 4-OH tamoxifen at stage P3 to permanently label Deiter's cells and pillar cells (SC-subtypes) with tdTomato (red-fluorescent protein). Explants were treated with and without JAG1-Fc peptide and were cultured in the presence of CHIR99021 and LY411575 to induce SC-to-HC conversion for 4 days. **b** High-power confocal images of SC-layer taken at the mid-apex of control and JAG1-Fc-peptide treated explants. Co-expression of tdTomato (red) and Atoh1-nGFP (green) labels SCs that converted into HCs. Scale bars=25 µm. **c** Quantification of newly formed HCs (tdTomato+; Atoh1-nGFP+) in (b) (n=3 animals per group). The two-tailed, paired Student's t test was used to calculate *P* value. **P ≤0.05.*

Supplementary Tables 1-3 are provided as separate files in spreadsheet format (excel file)

Supplementary Table 1: Gene expression Ctrl and *Jag1 KO*

Supplementary Table 2: DEG Ctrl versus *Jag1* KO

Supplementary Table 3: GO analysis DEG

**Supplementary Table 4. List of genotyping primers.**

| **Mouse line** | **Genotyping primers** | **Product size** |
|---|---|---|
| *Atoh1-nGFP*<br>*Lfng-GFP*<br>*P27-GFP* | EGFP1: CGA AGG CTA CGT CCA GGA GCG CAC<br>EGFP2: GCA CGG GGC CGT CGC CGA TGG GGG TGT | TG= 300bp |
| *Fgfr3icreER* | iCre-F: GAG GAC TAC CTC CTG TAC C<br>iCre-R: TGC CCA GAG TCA TCC TTG G | TG:600bp |
| *Jag1* $^{fl/fl}$ | Jag1-1: TGA ACT CAG GAC AGT GCT C<br>Jag1-2: GTT TCA GTG TCT GCC ATT GC<br>Jag1-3: ATA GGA GGC CAT GGA TGA CT | FL=500bp<br>WT=400bp<br>KO=330bp |
| *Notch1* $^{fl/fl}$ | Notch1-F: TGC CCT TTC CTT AAA AGT GG<br>Notch1-R: GCC TAC TCC GAC ACC CAA TA | TG:300bp<br>WT=250bp |
| *Notch2* $^{fl/fl}$ | Notch2-F: TAG GAA GCA GCT CAG CTC ACA<br>Notch2-R: ATA ACG CTA AAC GTG CAC TGG AG | MT=240bp<br>WT=202bp |
| *Notch3* $^{-/-}$ | Notch3-M-F: TCG CCT TCT ATC GCC TTC TTG<br>Notch3-M-R: GGT ACT GAG AAC CAA ACT CAG<br>Notch3-WT-F: CCA TGA GGA TGC TAT CTG TGA C<br>Notch3-WT-R: CAC ATT GGC ACA AGA ATG AGC C | MT=380bp<br>WT=287bp |
| *Pou4f3* $^{DTR/+}$ | Common: AAG AAG CAG GTG GGG GAG AG<br>Wild type Reverse: ATT GTT CTG GGC GAC ATG A<br>Mutant Reverse: CAG AAA GAG CTT CAG CAC CAC | WT:351bp<br>MT: 290bp |
| *R26 rtTA*M2* | MTR: GCG AAG AGT TTG TCC TCA ACC<br>F: AAA GTC GCT CTG AGT TGT TAT<br>WTR: GGA GCG GGA GAA ATG GAT ATG | WT=650bp<br>MT=340bp |

| | | |
|---|---|---|
| *TetO-cre*<br>*Emx2$^{cre/+}$*<br>*Sox2$^{CreER/+}$* | F: GCC TGC ATT ACC GGT CGA TGC AAC GA<br>R: GTG GCA GAT GGC GCG GCA ACA CCA TT | TG:700bp |

**Supplementary Table 5. List of qPCR primers.**

| Gene | Forward Primer | Reveres Primer |
|---|---|---|
| *Atoh1* | ATG CAC GGG CTG AAC CA | TCG TTG TTG AAG GAC GGG ATA |
| *Dkk3* | TGT GTA CAC TGC TGG CGG CG | GAG CTC TCC CTC CAC GGG CA |
| *Eif4ebp1* | AGC CAT TCC TGG GGT CAC TA | ATC ATT GCG TCC TAC GGC TG |
| *Emx2* | GAA TCC GCT TTG GCT TTC TG | GAC ACA AGT CCC GAG AGT TTC C |
| *Gfi1* | AGGAACGCAGCTTTGACTGT | TGAGATCCACCTTCCTCTGG |
| *Hes1* | GCT TCA GCG AGT GCA TGA AC | CGG TGT TAA CGC CCT CAC A |
| *Hey1* | CAC TGC AGG AGG GAA AGG TTA T | CCC CAA ACT CCG ATA GTC CAT |
| *HeyL* | GCG CAG AGG GAT CAT AGA GAA | TCG CAA TTC AGA AAG GCT ACT G |
| *Id1* | GAA CGT CCT GCT CTA CGA CAT G | TGG GCA CCA GCT CCT TGA |
| *Jag1* | TGT GCA AAC ATC ACT TTC ACC TTT | GCA AAT GTG TTC GGT GGT AAG AC |
| *Lfng* | ACT GCA CCA TTG GCT ACA TTG | GGC CGC TCC GGA TGA |
| *Minar2* | AGA ACA ACC CGT TGT ATG GTG A | GTC GAA GCC AGG AGT GTA CG |
| *Myo7a* | CCC CCT CTG AGA AGT TCG TTA A | TGT GTC CGA GTT CCG TTG AC |
| *Notch1* | CGT GGT CTT CAA GCG TGA TG | AGC TCT TCC TCG TGG CCA TA |
| *Notch2* | TCT ATC CCC CGT CGA TTC G | GAT GTG ATC ATG GGA GAG GAT GT |
| *Nupr1* | AAA AAG CCT GGC GCT GAG A | GGC CTA GGT CCT GCT TAC AAC |
| *Pou4f3* | GCA CCA TCT GCA GGT TCG A | CCG GCT TGA GAG CGA TCA T |
| *Rpl19* | GGT CTG GTT GGA TCC CAA | TGC CCG GGA ATG GAC AGT CA |
| *S100a1* | TGG ATG TCC AGA AGG ATG CA | CCG TTT TCA TCC AGT TCC TTC A |
| *Scx* | CTT CAC TGC GCT GCG CAC ACT | GCT CTC CGT GAC TCT TCA GTG |
| *Sox2* | CCA GCG CAT GGA CAG CTA | GCT GCT CCT GCA TCA TGC T |

**Supplementary Table 6.** Antibodies for Immunostaining

| Reagent type | Designation | Source | Identifiers | Additional information |
|---|---|---|---|---|
| antibody | Biotinylated donkey anti-goat | Jackson Immuno Research Lab | Cat.# 705-065-147 | 1:200 dilution |
| antibody | donkey anti-goat IgG (H+L) Alexa Fluor 546 | ThermoFisher | Cat.# A-11056 | 1:1000 dilution |
| antibody | donkey anti-rabbit IgG (H+L) Alexa Fluor 647 | ThermoFisher | Cat.# A-31573 | 1:1000 dilution |
| stain | Hoechst 33258 solution | Sigma-Aldrich | Cat.# 94403 | 1:3000 dilution |
| antibody | JAG1 goat polyclonal | Santa Cruz | Cat.# sc-6011 | 1:500 dilution |
| antibody | myosin VIIa rabbit polyclonal | Proteus Biosciences | Cat.# 25-6790 | 1:500 dilution |
| antibody | SOX2 goat polyclonal | Santa Cruz | Cat.# sc-17320 | 1:500 dilution |
| dye | Streptavidin, Alexa Fluor 405 | Life Technologies | Cat. #S32351 | 1:200 dilution |

**Supplementary Table 7.** Antibodies for Immunoblotting

| Reagent type | Designation | Source | Identifiers | Additional information |
|---|---|---|---|---|
| primary antibody | 4E-BP1 rabbit monoclonal | Cell Signaling | Cat.# 9644 | 1:2000 dilution |
| primary antibody | Akt rabbit monoclonal | Cell Signaling | Cat.# 4691 | 1:1000 dilution |
| primary antibody | Cleaved Notch1(Val1744) rabbit monoclonal | Cell Signaling | Cat.# 4147 | 1:1000 dilution |
| primary antibody | JAG1 goat polyclonal | Santa Cruz | Cat.# sc-6011 | 1:1000 dilution |
| primary antibody | Notch1 rabbit monoclonal | Cell Signaling | Cat.# 4830 | 1:1000 dilution |

| | | | | |
|---|---|---|---|---|
| primary antibody | Notch2<br>rabbit monoclonal | Cell Signaling | Cat.# 5732 | 1:1000 dilution |
| primary antibody | p-4E-BP1(Thr37/46)<br>rabbit monoclonal | Cell Signaling | Cat.# 2855 | 1:2000 dilution |
| primary antibody | p-Akt (Ser473)<br>rabbit monoclonal | Cell Signaling | Cat.# 4060 | 1:1000 dilution |
| primary antibody | p-S6(Ser240/244)<br>rabbit monoclonal | Cell Signaling | Cat.# 5364 | 1:1000 dilution |
| primary antibody | S6<br>Mouse monoclonal | Cell Signaling | Cat.#2317 | 1:1000 dilution |
| primary antibody | β-actin<br>mouse monoclonal | Santa Cruz | Cat.# 47778 | 1:500 dilution |
